# Fine chromatin-driven mechanism of transcription interference by antisense noncoding transcription

**DOI:** 10.1101/649434

**Authors:** Jatinder Kaur Gill, Andrea Maffioletti, Varinia García-Molinero, Françoise Stutz, Julien Soudet

**Affiliations:** Dept. of Cell Biology, University of Geneva, 1211 Geneva 4, Switzerland; Institut de Génétique Humaine, CNRS - Université de Montpellier, Montpellier, France

**Keywords:** Noncoding transcription, Antisense, Nucleosomes, Histone modifications, Nucleosome-depleted regions, Chromatin remodeler, RSC, Yeast

## Abstract

Eukaryotic genomes are almost entirely transcribed by RNA polymerase II (RNAPII). Consequently, the transcription of long noncoding RNAs (lncRNAs) often overlaps with coding gene promoters triggering potential gene repression through a poorly characterized mechanism of transcription interference. In this study, we propose a global model of chromatin-based transcription interference in *Saccharomyces cerevisiae* (*S. cerevisiae*). By using a noncoding transcription inducible strain, we analyzed the relationship between antisense elongation and coding sense repression, nucleosome occupancy and transcription-associated histone modifications using near-base pair resolution techniques. We show that antisense noncoding transcription leads to the deaceylation of a subpopulation of −1/+1 nucleosomes associated with increased H3K36 trimethylation (H3K36me3). Reduced acetylation results in decreased binding of the RSC chromatin remodeler at −1/+1 nucleosomes and subsequent sliding into the Nucleosome-Depleted Region (NDR) hindering Pre-Initiation Complex (PIC) association. Finally, we extend our model by showing that natural antisense noncoding transcription significantly represses around 20% of *S. cerevisiae* genes through this chromatin-based transcription interference mechanism.

**Highlights:**

- Induction of antisense noncoding transcription leads to −1/+1 nucleosome sliding that competes with sense transcription PIC deposition.
- Antisense induction leads to a subpopulation of H3K36me3 nucleosomes differently positioned compared to H3K18ac nucleosomes.
- RSC chromatin remodeler recruitment to −1/+1 nucleosomes is modulated by histone acetylation levels.
- 20% of *S. cerevisiae* genes are significantly repressed by this antisense-dependent chromatin-based transcription interference mechanism.

## Introduction

Recent techniques monitoring eukaryotic nascent transcription have revealed that the RNA polymerase II (RNAPII) landscape extends far beyond the sole transcription of mRNA (Core et al. 2008; Churchman and Weissman 2011; Mayer et al. 2015; Nojima et al. 2015). If 1-2% of the human genome is devoted to coding genes, more than 80% is transcribed into >200nt long noncoding RNAs (lncRNAs) (Djebali et al. 2012). Thus, around 200,000 lncRNAs originating from Nucleosome-Depleted Regions (NDRs) have been recently annotated across different human tissues and cell types (Kaikkonen and Adelman 2018). While their function is still under debate, it raises a new concept in which RNAPII transcribes nearly the whole genome as closely interleaved transcription units (Mellor et al. 2016). Consequently, transcription of many lncRNAs is reaching coding gene promoters, eventually leading to transcription interference, *i.e.* repression of the coding gene (Proudfoot 1986). So far, the precise molecular basis underlying transcription interference remains poorly characterized.

In *Saccharomyces cerevisiae* (*S. cerevisiae*), noncoding transcription often originates from a NDR, also referred to as bidirectional promoters, transcribing a coding gene in one orientation and a noncoding RNA in the other (Neil et al. 2009; Xu et al. 2009; Churchman and Weissman 2011; Jensen et al. 2013). Because the yeast genome is compact, a majority of lncRNAs appear as being antisense to coding genes. To avoid antisense transcription into sense paired promoters and to limit transcription interference, lncRNAs are usually subjected to early termination in a process dependent on the Nrd1-Nab3-Sen1 complex followed by degradation (Porrua and Libri 2015). Early termination of noncoding transcription is not strict and mostly depends on the number of Nrd1-Nab3 recognition motifs carried by the lncRNA (Schulz et al. 2013; Castelnuovo et al. 2014). This implies that some long noncoding RNAs will be cleared through early-termination while others will naturally extend into sense promoters, followed by export and degradation in the cytoplasm *via* Nonsense-Mediated Decay (NMD) (Malabat et al. 2015; Wery et al. 2018). Importantly, artificial loss of early termination through Nrd1 depletion leads to antisense elongation into paired sense promoters and to a subsequent transcription interference directly correlated with the levels of lncRNA accumulation over the promoters (Schulz et al. 2013). According to this view, high steady state levels of natural antisense transcription into gene promoters may anticorrelate with coding sense expression. However, this question is still open since some analyses have led to opposite conclusions (Murray et al. 2015; Brown et al. 2018; Nevers et al. 2018).

RNAPII transcription has an impact on chromatin structure (Rando and Winston 2012; Zentner and Henikoff 2013; Venkatesh and Workman 2015). First, the mean spacing between nucleosomes correlates with transcription frequency (Chereji et al. 2018). The more a gene is transcribed, the more the distance between nucleosomes decreases. Second, RNAPII elongation is associated with concomitant deposition of histone modifications. The Carboxy Terminal Domain (CTD) of RNAPII interacts with Histone Methyl Transferases (HMT) catalyzing the transfer of methyl groups to histone H3. Among them, trimethylation of H3 at lysine 36 (H3K36me3) by Set2 follows a specific pattern over coding genes, being enriched at mid-to end of genes. This modification is a recruitment platform for the Rpd3S Histone DeACetylase (HDAC) complex which can deacetylate H3 and H4 histones (Rundlett et al. 1996; Carrozza et al. 2005; Keogh et al. 2005). Accordingly, H3 and H4 acetylations anticorrelate with H3K36me3, being more enriched at −1 and +1 nucleosomes flanking the promoter NDRs (Weiner et al. 2015; Sadeh et al. 2016). Deletion of *SET2* or *RPD3* leads to intragenic cryptic transcription associated with an increase of histone acetylation along gene bodies (Li et al. 2003; Carrozza et al. 2005; Joshi and Struhl 2005; Venkatesh et al. 2012; Malabat et al. 2015; Venkatesh and Workman 2015; Kim et al. 2016). Thus, transcription-associated methylation of nucleosomes and subsequent deacetylation can be considered as a locking mechanism - limiting spurious transcription initiation events.

Interestingly, noncoding transcription-mediated transcription interference usually requires RNAPII-dependent nucleosome modifications. Indeed, loss of Set2 or HDACs leads to a rescue of coding gene expression when noncoding transcription elongates into promoter NDRs (Camblong et al. 2007; van Werven et al. 2012; Castelnuovo et al. 2014; Kim et al. 2016; du Mee et al. 2018; Nevers et al. 2018). Moreover, the more nascent transcription enters into sense promoters at steady-state, the more the NDR becomes narrow (Dai and Dai 2012; Murray et al. 2015). Altogether, these data suggest that antisense-mediated transcription interference probably occurs through a nucleosome-based mechanism. Indeed, nucleosome positioning at NDRs is crucial for the accessibility of the transcription Pre-Initiation Complex (PIC) to promoters, mainly through the recruitment of the TATA-Binding Protein (TBP, Spt15 in *S. cerevisiae*) at TATA (or TATA-like) Binding Sites (TBSs) (Rhee and Pugh 2012; Kubik et al. 2018). Importantly, genome-wide NDR opening and subsequent ability to recruit the PIC mainly depends on the RSC ATP-dependent chromatin remodeler (Badis et al. 2008; Hartley and Madhani 2009; Kubik et al. 2018; Klein-Brill et al. 2019).

In this study, we aim at defining a comprehensive mechanism of antisense-mediated transcription interference through a variety of genome-wide approaches. We use a strategy in which antisense noncoding early termination can be turned off, leading to inducible transcription interference of more than 200 genes. Systematic analyzes of transcription initiation, NDR chromatin structure and transcription-associated histone modifications reveal the choreography of chromatin-related events associated with lncRNA-induced transcription interference. We then validate our model by defining to which extent the *S. cerevisiae* genome is influenced by this chromatin-based transcription interference at steady state.

## Results

### Induction of antisense noncoding transcription into paired sense promoters decreases PIC binding

To investigate the mechanism of transcription interference by antisense noncoding transcription, we first defined the list of coding genes that are repressed upon abrogation of RNAPII early termination. To do so, we performed RNA-seq of the Nrd1-Anchor Away (AA) strain in which Nrd1 can be artificially depleted from the nucleus upon Rapamycin (Rap) addition (Fig. 1A) (Haruki et al. 2008; Schulz et al. 2013). We then classified the genes into three categories, i) the Antisense-Mediated Repressed Genes (AMRG, 217 genes), showing at least 2-fold increase in antisense and >20% sense repression, ii) the Non-Responsive Genes (NRG, 469 genes), with a minimum 2-fold increase in antisense but less than 20% decrease in sense expression and, iii) the Others (4089 genes), showing less than 2-fold increase in antisense (Fig. 1A, B). Taking these 3 groups, we observe a prominent correlation between the fold-increase of antisense levels over the paired sense promoter and the fold-decrease of sense transcription as already described (Fig. 1C) (Schulz et al. 2013). AMRG are significantly more repressed than the Others while the NRG present a mild phenotype. To directly assess nascent transcription, we took advantage of published datasets monitoring RNAPII PAR-CLIP in an Nrd1-AA strain (Schaughency et al. 2014). The same trend as obtained with the RNA-seq is observed, although with a lower amplitude, implying that sense repression may occur at the nascent transcription level (Fig. 1D) (Schaughency et al. 2014).

**Figure 1.**
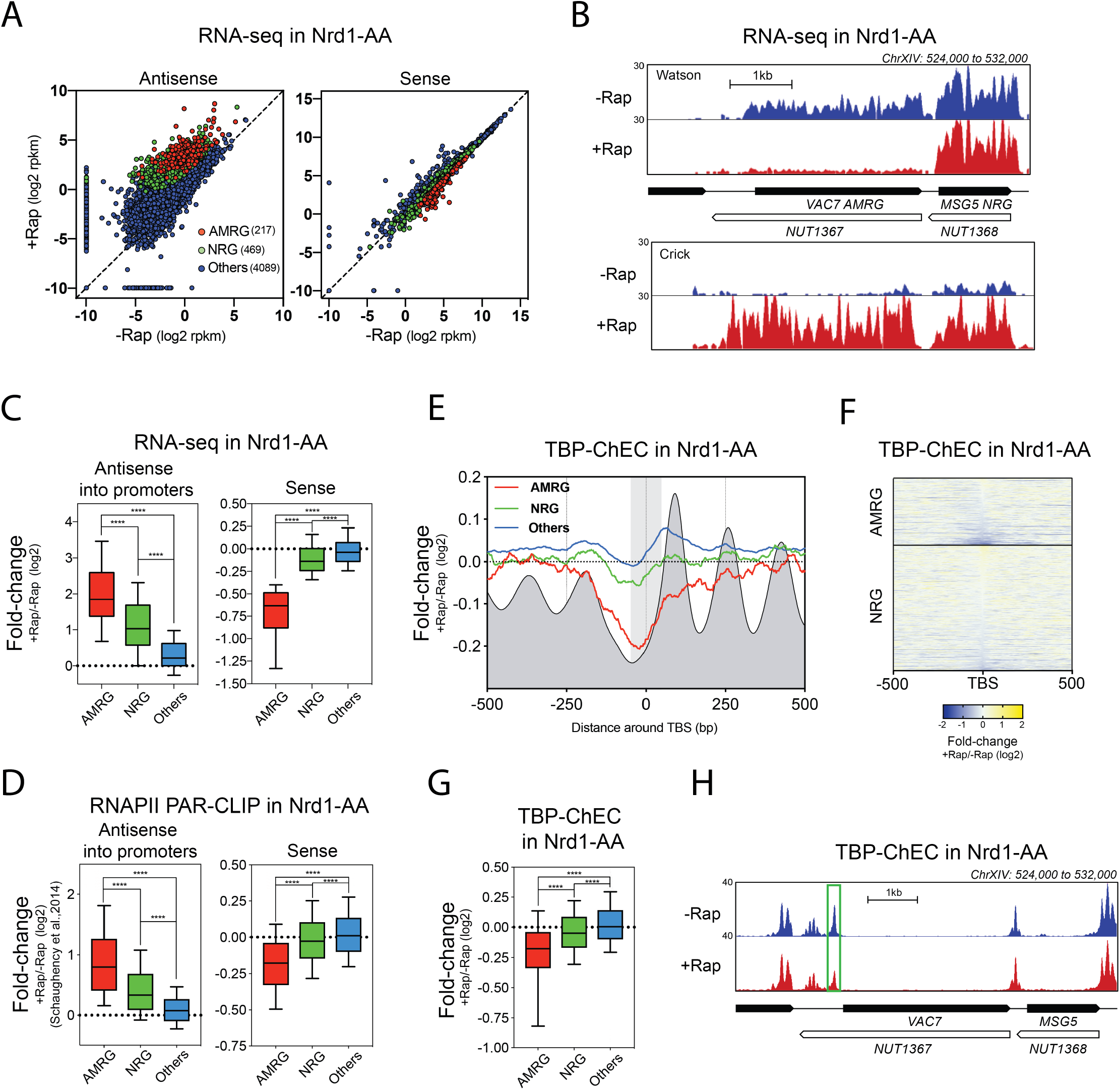
Antisense-mediated transcription interference decreases PIC binding. **(A)** Scatter dot-plot of the Nrd1-AA RNA-seq. Nrd1-AA cells were treated or not with Rapamycin (Rap) for 1h before RNA extraction. Results are represented in log2 of Reads Per Kilobase of transcript per Million mapped reads (rpkm) in both antisense and sense orientation. DEseq was used to define the different gene classes (see Method details). **(B)** Snapshot of the Nrd1-AA RNA-seq depicting a locus containing both an AMRG (*VAC7*) and an NRG (*MSG5*). Antisense induction in +Rap leads to the elongation of *NUT1367* noncoding RNA into the *VAC7* promoter and to the subsequent transcription interference of the *VAC7* transcript. **(C)** Box-plots showing the (+Rap/-Rap) fold-change for the RNA-seq in the Nrd1-AA strain. Fold-change is calculated based on the coverage in the −100bp to Transcription Start Site (TSS) region for antisense and over the whole transcript for the sense. **(D)** Box-plots indicating (+Rap/-Rap) fold-change for the RNAPII PAR-CLIP in an Nrd1-AA strain. Data were taken from (Schaughency et al., 2014). Antisense transcription was measured over the −100bp to TSS of the paired sense, and sense transcription was examined over 100bp upstream of the polyA site. **(E)** Metagene analysis of TBP-ChEC induced for 30sec in an Nrd1-AA strain treated or not with Rap for 1h. Colored curves represent (+Rap/-Rap) fold-change of the TBP-ChEC for the different classes of genes. The grey profile depicts the position of nucleosomes as obtained with MNase-seq (Fig. 2A) for the average gene. The grey box represents the 50bp area around the TBS. The center of 0-120bp paired-end fragments is represented for the plots. **(F)** Heatmap of (+Rap/-Rap) fold-change of TBP-ChEC for the AMRG and NRG. **(G)** Box-plots indicating the (+Rap/-Rap) fold-change of TBP-ChEC in the Nrd1-AA strain as measured 50bp around the TBS. **(H)** Snapshot depicting the AMRG-class coding gene *VAC7* repressed in +Rap by induction of *NUT1367* antisense transcription. The rectangle highlights the PIC involved in *VAC7* transcription.

Since sense repression may be transcriptional and related to the extension of antisense into the promoters, we investigated whether sense transcription initiation is affected upon antisense induction. To address this question, we performed Chromatin-Endogenous Cleavage (ChEC) of the TATA-Binding Protein (TBP, Spt15 in *S. cerevisiae*) in the Nrd1-AA strain treated or not with Rap in order to visualize PIC recruitment (Zentner et al. 2015). By ChEC, we observed the expected distribution of the PIC as peaks around the TATA (-like) Binding Sites (TBS) (Supplemental Fig. S1A) (Rhee and Pugh 2012). Interestingly, AMRG present a stronger decrease in PIC binding upon antisense induction as compared to the Others, while NRG again exhibit a milder phenotype (Fig. 1E-H; Supplemental Fig. 1B, C). The almost perfectly overlapping PIC peaks in -Rap/+Rap at the Others demonstrate the precision of the measurement and validate the experimental accuracy (Supplemental Fig. 1B).

Hence, as inferred by lower binding of the PIC at coding gene promoters, sense repression by antisense noncoding transcription occurs at the transcription initiation step.

### Antisense induction into promoters increases nucleosome occupancy over the TBS through −1/+1 sliding

As mentioned above, antisense-mediated transcription interference involves a chromatin-based mechanism. An interesting scenario would be that nucleosomes repositioned over promoters upon antisense induction may compete with PIC binding. We investigated this possibility at near base-pair resolution by monitoring nucleosome positioning around TBSs by paired-end Micrococcal Nuclease-sequencing (MNase-seq) upon antisense elongation in +Rap (Fig. 2A). To visualize the nucleosomes overlapping with the TBSs, we plotted nucleosome dyads and quantified the occupancy over a 100bp TBS-centered region. We detect a highly significant increase of nucleosome occupancy at the AMRG TBS as compared to the NRG and the Others (Fig. 2A, B, I; Supplemental Fig. S2A, B). To rule out the possibility that this measurement may be artifactual due to near-background values, we performed a complementary technique, Assay for Transposase-Accessible Chromatin-sequencing (ATAC-seq), to convert the “valley” signal into a “peak” signal, making this measurement more accurate. Similar to the MNase-seq, antisense induction into AMRG promoters significantly reduces the accessibility of the NDR (Fig. 2C, D and I). Even <150bp NDRs, in which an additional nucleosome cannot fit, show this increase in nucleosome occupancy suggesting −1 and/or +1 sliding rather than incorporation of new nucleosomes (Fig. 2E).

**Figure 2.**
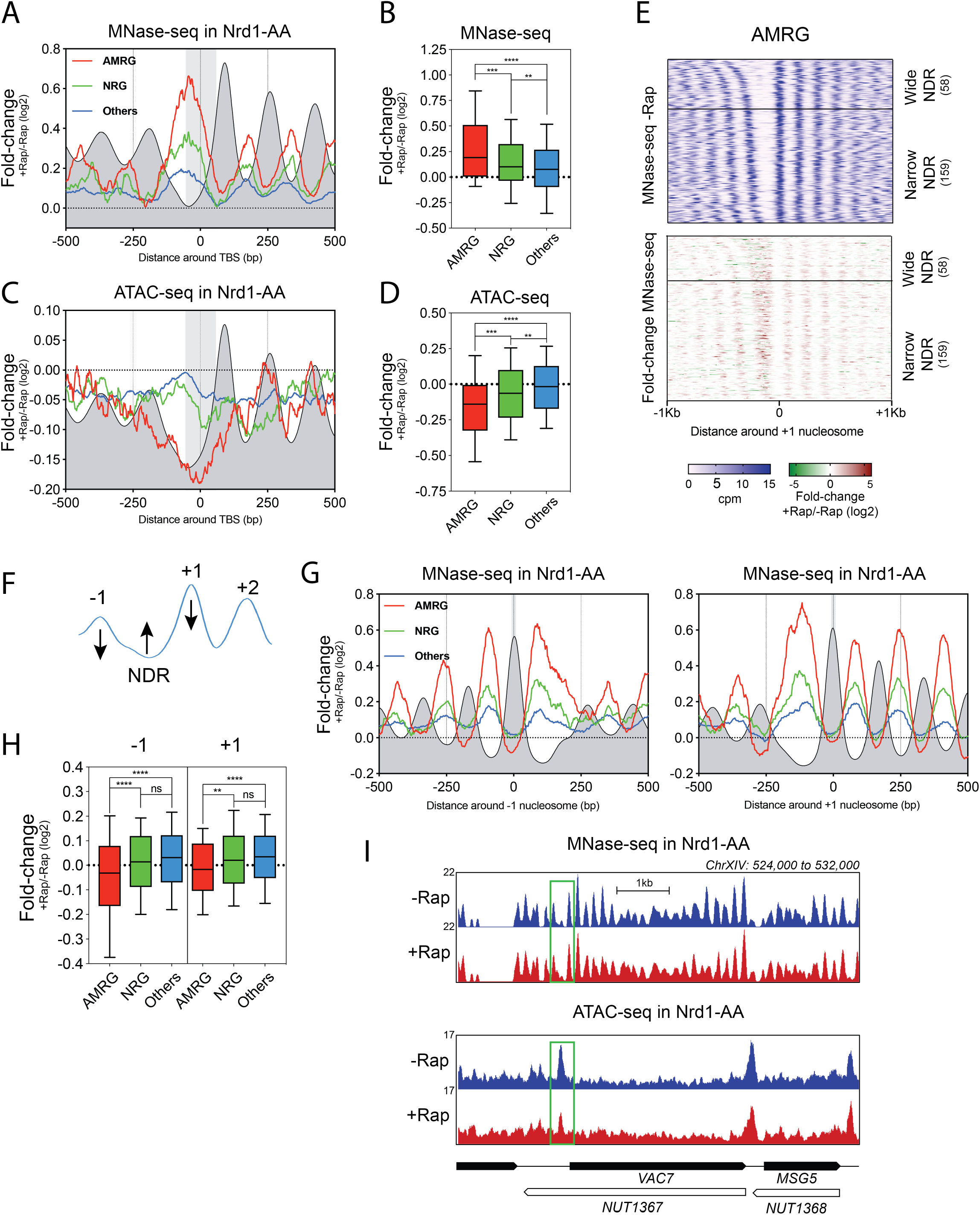
Antisense induction leads to paired sense promoter closing. **(A)** Metagene analysis of MNase-seq obtained in an Nrd1-AA strain treated or not with Rap for 1h. Colored curves represent the (+Rap/-Rap) fold-change of the MNase-seq for the different classes of genes. The grey profile depicts the position of nucleosomes as obtained for the average gene. The grey box represents the 50bp area around the TBS. The center of 120-200bp paired-end fragments is represented for the plots. **(B)** Box-plots indicating the (+Rap/-Rap) fold-change of dyads occupancy. Statistics are generated in the 50bp-TBS centered area. **(C)** Metagene analyses of ATAC-seq in an Nrd1-AA strain treated or not for 1h with Rap and represented as in (A). **(D)** Box-plots indicating the (+Rap/-Rap) fold-change in NDR accessibility as measured with the ATAC-seq. Statistics are generated as in (B). **(E)** Heatmaps centered on the +1 nucleosome of MNase-seq signal in -Rap (top) and MNase-seq (+Rap/-Rap) fold-change (bottom) for the AMRG. NDRs were ranked according to their length to discriminate between “Wide” NDRs, in which additional nucleosomes can virtually accommodate, and “Narrow” NDRs that cannot fit a 150bp DNA-covered nucleosome. **(F)** Cartoon depicting that gain of nucleosomes around the TBSs through sliding should be accompanied by −1 and /or +1 decrease in dyads occupancy peaks. **(G)** Metagene analysis of MNase-seq as in (A) but centered on the −1 and +1 nucleosomes respectively. The grey box represents the 10bp area around −1 and +1 peaks. **(H)** Box-plots representing the dyad occupancy (+Rap/-Rap) fold-change in a 10bp area over the −1 and +1 peaks respectively. **(I)** Snapshot of both MNase-seq and ATAC-seq profiles at the AMRG *VAC7*. The NDR from which *VAC7* transcription is initiated is highlighted by a rectangle.

If −1 and/or +1 nucleosomes were sliding, one would expect a decrease of the occupancy when centering the analysis on their dyads (Fig. 2F). This is what we observe for the −1 and +1 of AMRG upon antisense induction (Fig. 2G, H; Supplemental Fig. S2A). This AMRG specific decrease is subtle in terms of fold-change but nicely anticorrelates with the fold-increase gained over TBSs. Importantly, when performing Chromatin ImmunoPrecipitation (ChIP) of H3 with sonicated extracts (with around 300 bp-resolution) at AMRG promoters, we do not detect any change in H3 content upon antisense induction, justifying the need of near base-pair resolution to visualize antisense-mediated chromatin changes (Supplemental Fig. S2C).

Importantly, PIC depletion does not lead to −1/+1 sliding at AMRG (Supplemental Fig. S1D, E). This strongly suggests that decreased PIC binding is a consequence of nucleosome repositioning rather than the opposite.

Altogether, these data show that antisense induction into AMRG promoters leads to the repositioning of a subpopulation of −1/+1 nucleosomes over the TBS thereby competing with the recruitment of the PIC. Of note, increase in nucleosome occupancy at AMRG is not restricted to the NDR as it also appears to increase in between +1/+2, +2/+3 and so on (Fig. 2G, I). Thus, antisense induction not only leads to promoter rearrangement but also to phasing changes of the gene body nucleosomal array.

### H3K36me3 levels increase over AMRG TBSs upon antisense induction

Since H3K36me3 by the Set2 HMT is known to be involved in ncRNA-mediated transcription interference at several individual genes (van Werven et al. 2012; Kim et al. 2016; du Mee et al. 2018; Nevers et al. 2018), we analyzed this modification at nucleosome-resolution using MNase-ChIP-seq (Weiner et al. 2015). Upon antisense elongation, we detect an increase in H3K36me3 over TBSs but not when measuring at −1 and +1 nucleosome dyads (Fig. 3A, B, G; Supplemental Fig. S3A, B). This increase in H3K36me3 is not biased towards large NDRs suggesting that this modification is linked to the sliding event and corresponds to a *de novo* modification induced by noncoding transcription entering into the NDR (Fig. 3E). In agreement, H3K36me3 ChIP at individual gene promoters indicates an increase of this modification at AMRG with longer incubation times in Rap, strengthening the model that antisense elongation into promoters induces *de novo* H3K36me3 deposition (Fig. 3F).

**Figure 3.**
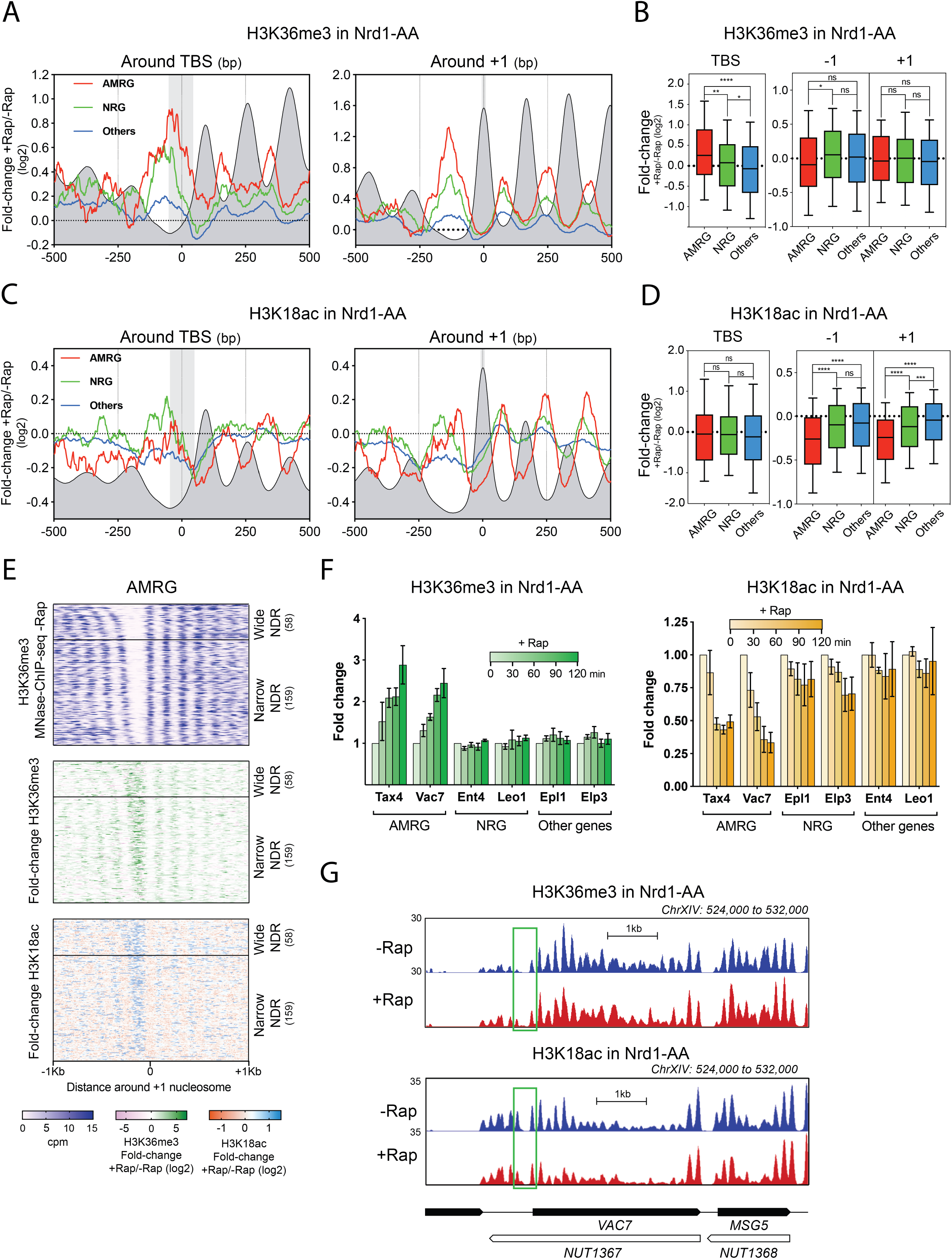
Antisense elongation into AMRG promoters leads to increased H3K36me3 levels over TBS and decreased H3K18ac levels at −1/+1 nucleosomes. **(A)** Metagene profiles of H3K36me3 MNase-ChIP-seq centered on TBSs and +1 nucleosome dyad obtained in an Nrd1-AA strain treated or not for 1h with Rapamycin. Colored curves represent (+Rap/-Rap) fold-change of the H3K36me3 modification for the different classes of genes. The grey profile depicts the position of H3K36me3 nucleosomes as obtained for the average gene in the absence of Rapamycin. The grey boxes represents the 50bp area around the TBS and the 10bp-centered +1 peak area. The center of 120-200bp paired-end fragments is represented for the plots. **(B)** Box-plots indicating the (+Rap/-Rap) fold-change in H3K36me3 levels in the 50bp TBS- and 10bp −1/+1-centered areas. **(C)** Metagene profiles as in (A) but for the H3K18ac levels. The grey profile represents the position of H3K18ac nucleosomes for the average gene in the absence of Rapamycin. **(D)** Box-plots as in (B) for the H3K18ac modification. **(E)** Heatmaps of H3K36me3 and H3K18ac fold-change centered on the +1 nucleosome dyad at AMRG. **(F)** ChIP of H3K36me3 and H3K18ac histone modifications at gene promoters of AMRG, NRG and Other genes. ChIPs were performed at 0, 30, 60, 90 and 120min after rapamycin addition. Immunoprecipitated promoter NDRs were normalized to immunoprecipitated *SPT1*5 ORF after qPCR amplification. Primers were designed to target each promoter NDR. The fold-change was artificially set to 1 for each gene in the -Rap condition. Error bars represent the Standard Error of the Mean (SEM) for a set of 3 independent experiments. **(G)** Snapshot of H3K36me3 and H3K18ac MNase-ChIP-seq levels at the AMRG *VAC7*. The NDR of *VAC7* is highlighted by a rectangle.

Thus, the −1/+1 nucleosome repositioning correlates with the appearance of H3K36me3 over the TBS induced by antisense transcription, suggesting that newly-modified H3K36me3 nucleosomes undergo a sliding event.

### H3K18ac levels decrease at −1/+1 AMRG nucleosomes upon antisense induction

Histone deacetylases have previously been shown to be involved in antisense-mediated transcription interference (Camblong et al. 2007; Castelnuovo et al. 2013). Hence, we analyzed the H3K18ac modification landscape by MNase-ChIP-seq in the Nrd1-AA strain. In contrast to H3K36me3, H3K18ac levels specifically decrease at −1/+1 nucleosomes but do not change over the TBSs following antisense induction (Fig. 3C-E, G; Supplemental Fig. S3C, D).

Thus, H3K36me3 and H3K18ac present an opposite pattern of changes. When H3K36me3 levels increase over the TBS, H3K18ac levels decrease at −1/+1 nucleosomes. Since H3K36me3 is a platform to recruit a histone deacetylase and that H3K36me3 and H3K18ac are largely anti-correlated, it is tempting to speculate that antisense elongation into promoters may lead to *de novo* H3K36me3 of the −1/+1 nucleosomes triggering deacetylation and subsequent sliding (Carrozza et al. 2005; Keogh et al. 2005; Weiner et al. 2015; Sadeh et al. 2016).

### Induced antisense transcription reveals different positioning of H3K36me3- and H3K18ac-containing nucleosomes

The different behaviors of H3K36me3 and H3K18ac at AMRG NDRs prompt us to propose that H3K36me3- and H3K18ac-containing nucleosomes may be differentially repositioned upon antisense induction and that MNase-seq recapitulates the sum of these changes. We reasoned that since antisense is expressed all along the gene bodies upon Nrd1 anchor-away, the same trend may be observed within the nucleosomal array of gene bodies. We therefore performed a metagene analysis of the nucleosomes +1 to +8 for the different gene classes using both MNase-seq as well as H3K36me3 and H3K18ac MNase-ChIP-seq data (Fig. 4A). This approach increases the robusteness of our measurements since 1’223, 2’981 and 19’899 nucleosomes are taken into account in our analysis for AMRG, NRG and Others, respectively. We find that for AMRG, H3K36me3 levels do not significantly change at the peak but increase upstream and downstream of the peak, while H3K18ac levels decrease at the peak without any change on either side (Fig. 4A and B). The MNase-seq appears as the sum of these two events since nucleosome occupancy significantly decreases at the peak and increases upstream of the the peak.

**Figure 4.**
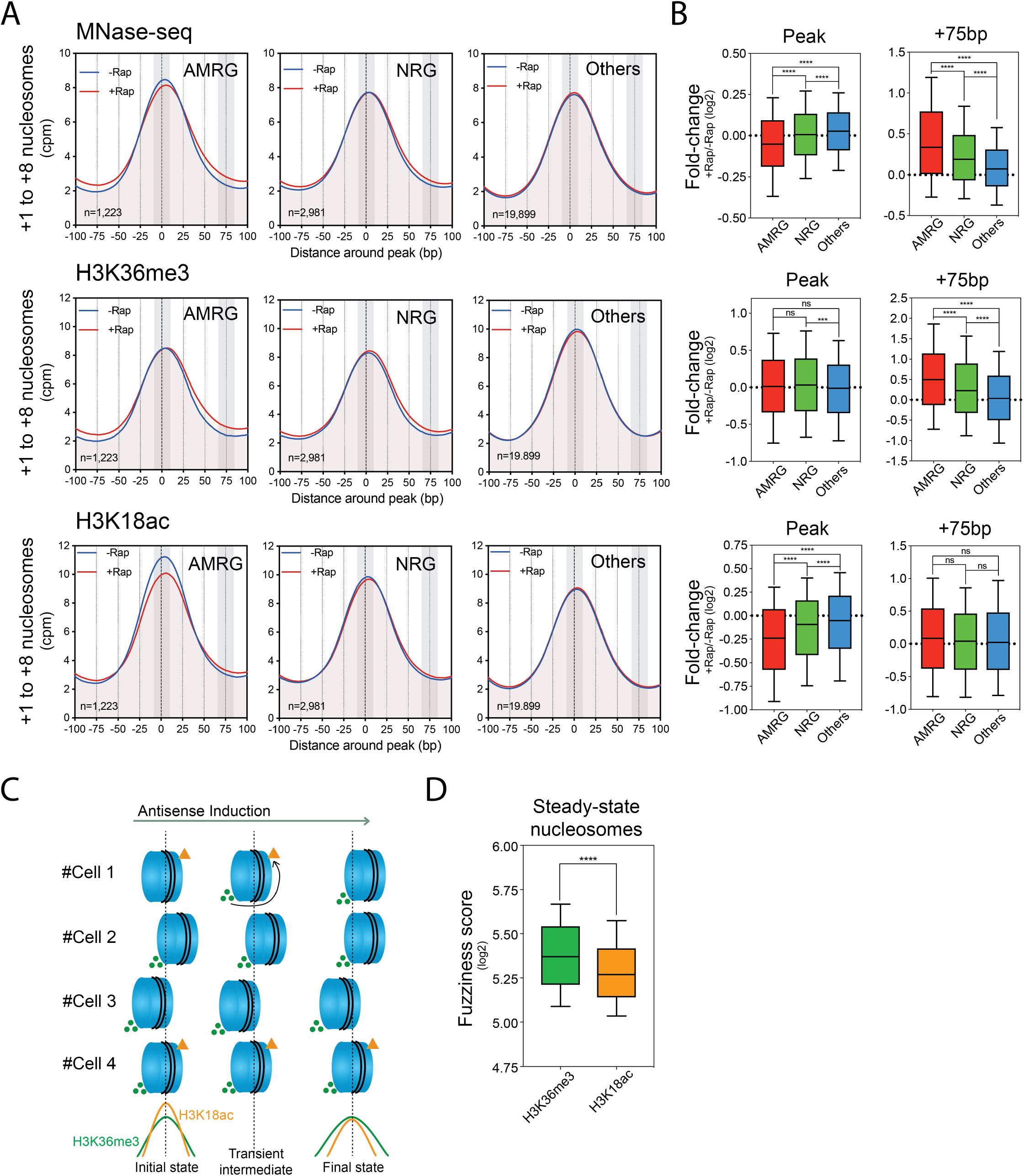
Different positioning of *de novo* H3K36me3-containing nucleosomes and H3K18ac-containing nucleosomes. **(A)** Metagene profiles of MNase-seq, H3K36me3 and H3K18ac MNase-ChIP-seq around an average nucleosome peak (+1 to +8 nucleosome peaks) upon antisense induction. Nucleosomes were averaged according to the directionality of the sense coding gene. The grey boxes represent the 10bp area around the nucleosome peak and the 10bp area located at +75bp, downstream of the nucleosome peak. **(B)** Box-plots indicating the (+Rap/-Rap) fold-change in MNase-seq, H3K36me3 and H3K18ac levels in the 10bp TBS- and 10bp +75bp-centered areas. **(C)** Model recapitulating the observed differences in positioning. Nucleosomes depicted represent the same nucleosome in 4 different cells upon antisense induction. **(D)** Fuzziness score of H3K36me3 and H3K18ac nucleosomes at steady-state (-Rap).

Thus, our data converge towards a model where antisense induction leads to *de novo* H3K36me3, associated H3K18 deacetylation and subsequent repositioning of nucleosomes (Fig. 4C). It implies that H3K18ac-containing nucleosomes may be restricted to the peaks while H3K36me3-containing nucleosomes may occupy a larger territory by generating several subpopulations: those at the peak that are both H3K36me3 and H3K18ac, and those around the peak that are only H3K36me3. Accordingly, at steady state, H3K36me3 nucleosomes are less well-positionned than H3K18ac nucleosomes (Fig. 4D).

### Decreased RSC binding at −1/+1 AMRG nucleosomes upon antisense induction

Considering the global role of the RSC chromatin remodeler in promoter NDR maintenance (Badis et al. 2008; Hartley and Madhani 2009; Yen et al. 2012; Kubik et al. 2018; Klein-Brill et al. 2019), we asked whether nucleosome sliding was due to loss of RSC interaction with the NDR-flanking nucleosomes. This possibility is even more appealing when considering that the RSC complex interacts with acetylated histones *in vitro* through its multiple bromodomains (Kasten et al. 2004; Chatterjee et al. 2011).

Consistently, when the recruitment of the Sth1 catalytic subunit of the RSC complex was examined by ChEC-seq in the Nrd1-AA strain, a specific decrease of RSC interaction with −1 and +1 AMRG nucleosomes was detected in the presence of Rap (Fig. 5A-C; Supplemental Fig. S4A, B). These observations suggest a link between the decreased acetylation of NDR-flanking nucleosomes and the loss of interaction with the essential RSC chromatin remodeler.

**Figure 5.**
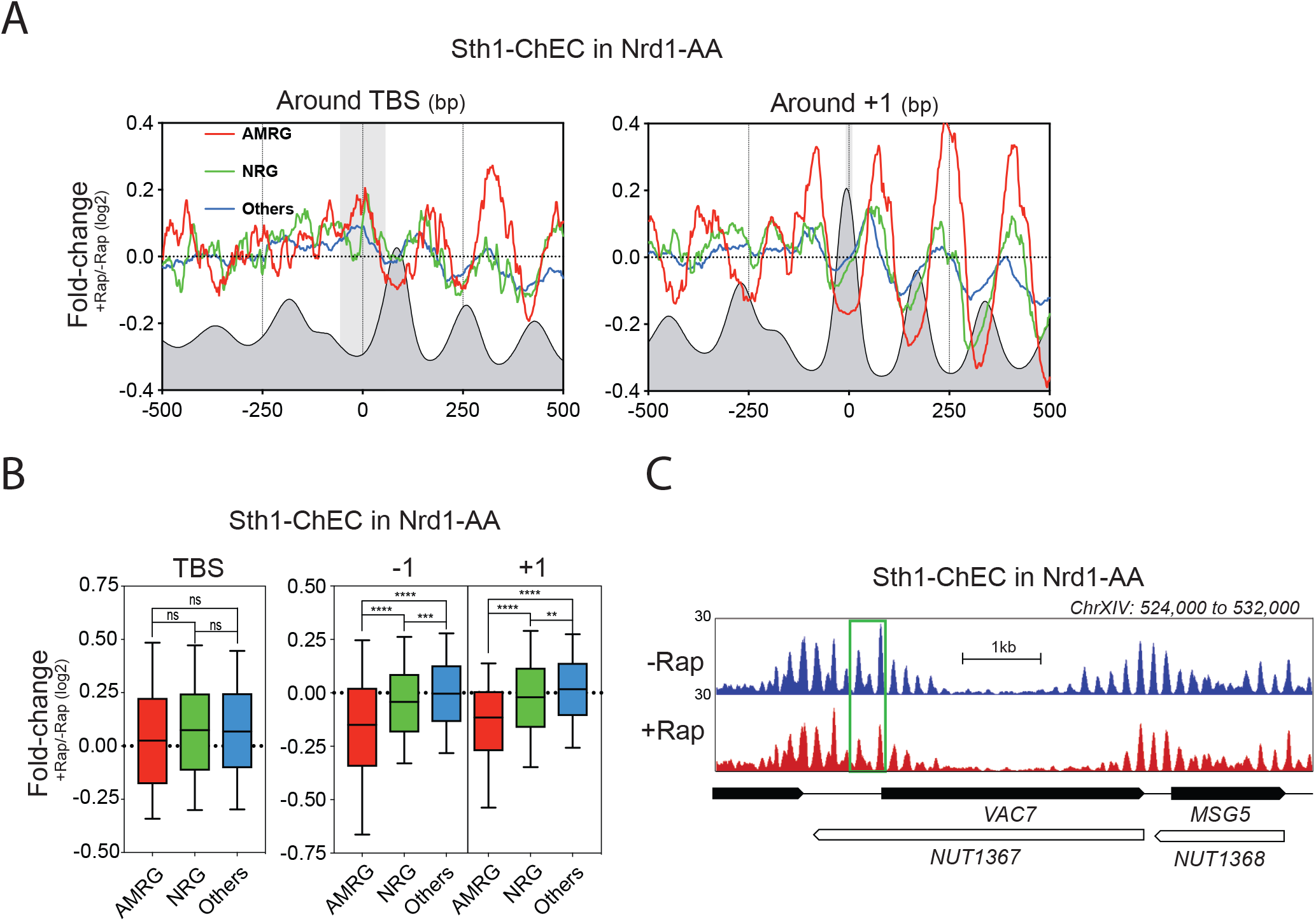
RSC interaction with −1/+1 nucleosomes decreases upon antisense induction. **(A)** Metagene profiles of Sth1-ChEC in an Nrd1-AA strain treated or not for 1h with Rap around the TBS and +1 nucleosome. Colored curves represent (+Rap/-Rap) fold-change of the Sth1-ChEC for the different classes of genes. The grey profile depicts the position of Sth1-ChEC as obtained for the average gene in the absence of Rapamycin. The grey boxes represent the 50bp area around the TBS and the 10bp-centered +1 peak area. The center of 120-200bp paired-end fragments is represented for the plots. **(B)** Box-plots indicating the (+Rap/-Rap) fold-change in Sth1-ChEC levels in the 50bp TBS- and 10bp--1/+1-centered areas. **(C)** Snapshot of Sth1-ChEC levels at the AMRG *VAC7*. The −1/+1 nucleosomes are highlighted by a rectangle.

### Loss of Rpd3 HDAC partially rescues transcription interference-associated phenotypes

If decreased −1/+1 acetylation directly affects RSC recruitment, a mutant in which the deacetylation step is defective may rescue RSC binding. This question was addressed by deletion of *RPD3* in the Nrd1-AA background. Rpd3 is the histone deacetylase component of the Rpd3S/L complex that maintains H3K18ac levels low in the cell (Rundlett et al. 1996).

We first examined the fold-change in H3K18ac levels by ChIP upon antisense induction at selected AMRG promoters in presence or absence of Rpd3 (Supplemental Fig. S5A). As expected, upon Rap addition, *RPD*3 deletion leads to limited deacetylation of AMRG NDR-flanking nucleosomes as compared to the *WT* strain.

We then plotted the +Rap/-Rap fold-change of RSC binding at the +1 nucleosome, as assayed by Sth1 ChEC-seq (Fig. 6A). When comparing to *WT* cells, we observe a partial rescue of RSC binding in *rpd3Δ* at AMRG. As RSC is essential for the maintenance of NDR opening (Badis et al. 2008; Hartley and Madhani 2009; Kubik et al. 2018; Klein-Brill et al. 2019), the increased retention of RSC at AMRG −1/+1 nucleosomes in *rpd3Δ* may inhibit the nucleosome sliding as compared to *WT* cells. Indeed, as revealed by ATAC-seq, the increase in nucleosome occupancy over the TBS observed in *WT* cells upon antisense induction is partially abrogated in *rpd3Δ* (Fig. 6B). Consequently, loss of PIC binding and transcription interference at AMRG observed in *WT* are also alleviated in *rpd3Δ* (Fig. 6C, D). Importantly, the level of antisense accumulation in AMRG is not globally affected in Nrd1-AA *rpd3Δ* as compared to Nrd1-AA cells, indicating that the observed rescue is not an indirect effect of decreased antisense transcription (Supplemental Fig. S5B).

**Figure 6.**
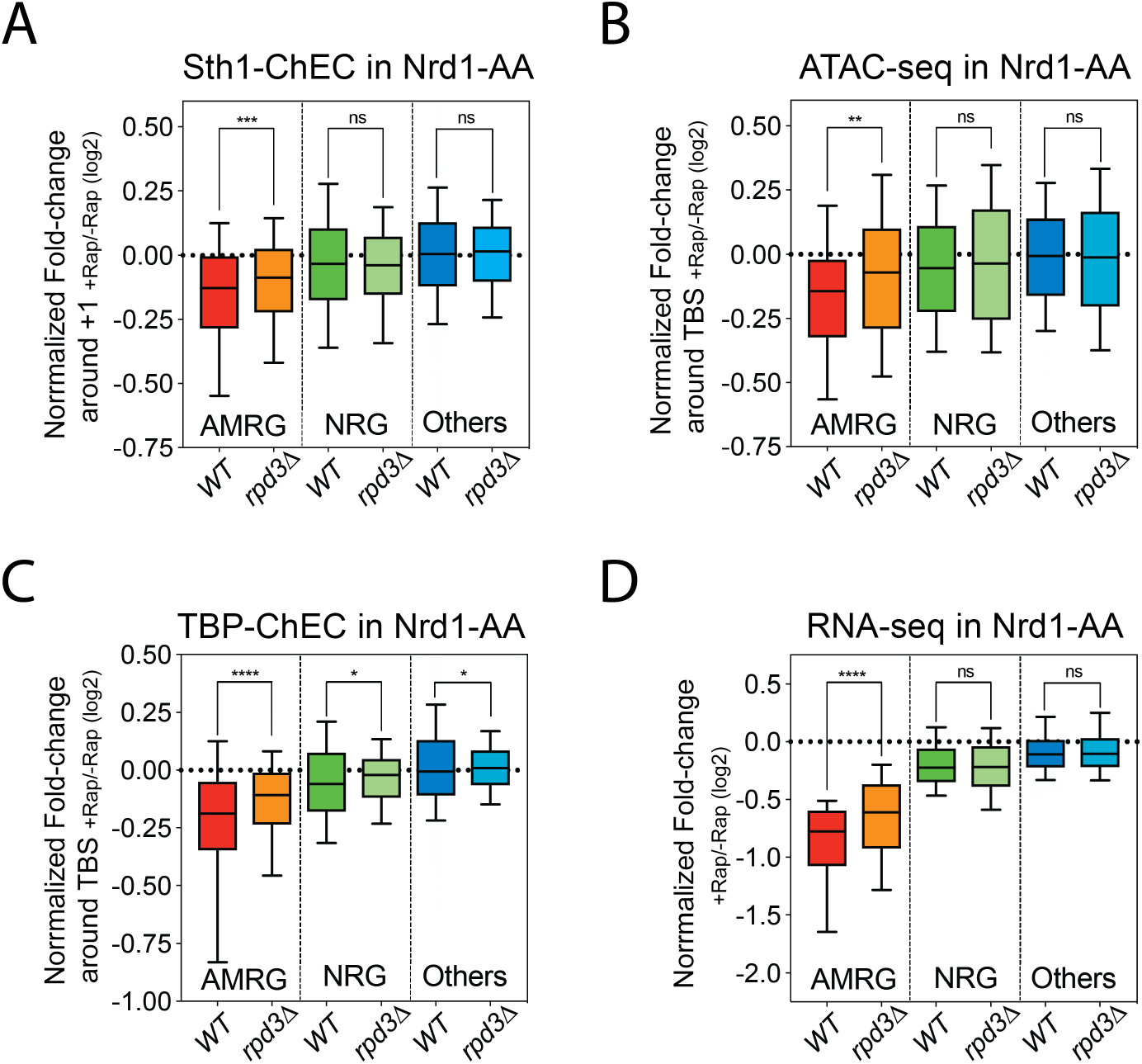
Absence of Rpd3 HDAC partially rescues antisense-mediated gene repression phenotypes. **(A) (B) (C) (D)** Box-plots indicating the normalized (+Rap/-Rap) fold-change for Sth1 ChEC-seq, ATAC-seq, TBP ChEC-seq and RNA-seq in an Nrd1-AA strain as compared with Nrd1-AA *rpd3Δ* cells. Plots for the Nrd1-AA strain are already presented in Figures 5B, 2B, 1G and 1C without the strain normalization, for which calculation is described in Method details.

Thus, the acetylation level of −1/+1 nucleosomes, through its ability to recruit the RSC chromatin remodeler, appears as a central regulator of the transcription interference mechanism. Consequently, acetylation influences the accessibility of the PIC to the NDR and hence gene expression.

### Model for antisense-mediated transcription interference in an inducible system and its genome-wide generalization at steady-state condition

Altogether, our results suggest the following model (Fig. 7A). Under normal conditions (-Rap), noncoding transcription early termination prevents entry of antisense transcription into the NDR of promoters maintaining them open and favoring gene expression. When Nrd1 is anchored away, antisense elongation extends into the paired promoter NDR, resulting in H3K36me3 of the −1 and +1 nucleosomes, which are subsequently deacetylated through a process involving the Rpd3 HDAC. Loss of acetylation leads to decreased RSC recruitment and subsequent sliding of the −1/+1 nucleosomes towards the TBS. These events result in a steric hindrance for PIC binding and, consequently, in gene repression.

**Figure 7.**
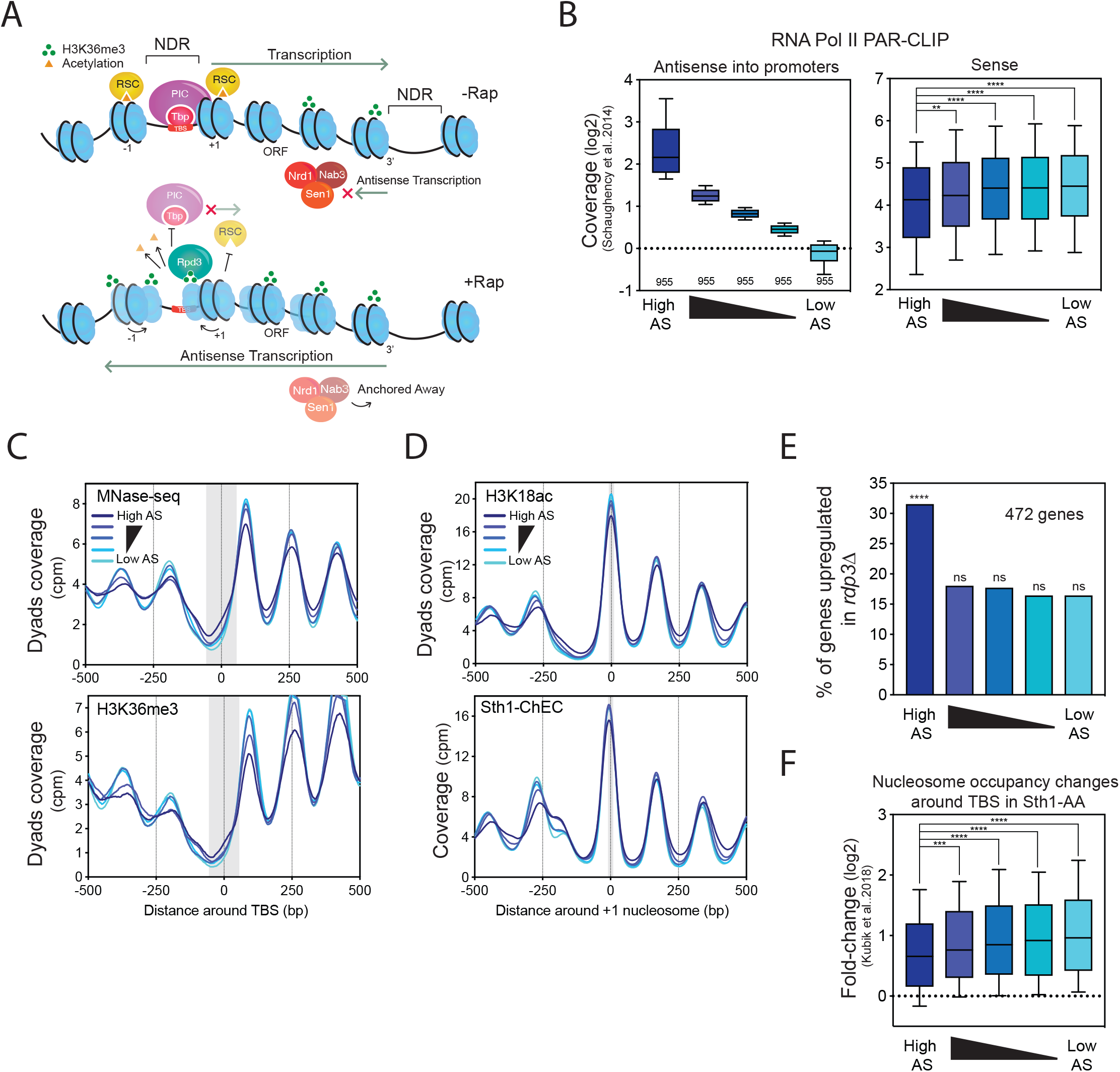
Model of induced chromatin-driven transcription interference and its generalization in the steady-state genome. **(A)** Model of antisense-mediated transcription interference through promoter chromatin regulation. For a description of the model, see the Results section. **(B)** Left panel: Box-plots defining the five quintiles according to their natural level of nascent antisense transcription into gene promoters (−100 to TSS area). Each quintile contains 955 genes. RNA PolII PAR-CLIP data were taken from (Schaughency et al., 2014). Right panel: Box-plots depicting levels of nascent coding sense transcription in an area of 100bp upstream of the polyA site. **(C)** Aggregate plot of nucleosome occupancy and H3K36me3 levels around the TBS with respect to the different quintiles. **(D)** Metagene analysis of H3K18ac (top) and RSC binding (bottom) levels around +1 nucleosome according to the different quintiles. **(E)** Percentage of genes from each quintile upregulated in a *rpd3Δ* strain (>2fold). A total of 472 genes are upregulated in the *rpd3Δ* strain. Significance was defined according to a hypergeometrical test. **(F)** Box-plot of nucleosome occupancy fold-change in a 50bp-TBS centered region upon anchor-away of Sth1 for 1h. Data were retrieved from (Kubik et al., 2018).

Our model is limited to only 4.5% of the genes in this non-physiological system to induce antisense elongation. Since antisense transcription can naturally extend into promoter NDRs depending on the strength of the noncoding transcription early termination process, we investigated whether our chromatin-based transcription interference model could be generalized under steady-state conditions. We ranked the coding genes into five quintiles according to the natural levels of nascent antisense transcription over their promoters (Fig. 7B). The genes of the first quintile, presenting the highest levels of antisense transcription into promoters, are significantly less expressed than the ones of the other quintiles as assessed by RNA-seq and using nascent transcription data (Fig. 7B; Supplemental Fig. S6A, B). As predicted by our model, the first quintile shows increased nucleosome occupancy and increased H3K36me3 levels over the TBS (Fig. 7C). Accordingly, the levels of H3K18ac and RSC binding at the +1 nucleosome appear as reduced in the first quintile compared to the others (Fig. 7D). Consistent with our model, in which H3K36me3-mediated deacetylation of NDR-flanking nucleosomes is involved in natural antisense-mediated transcription interference, the genes up-regulated in the absence of Rpd3 HDAC are significantly enriched in the first quintile (Fig. 7E). Lastly, we analyzed the sensitivity of the different quintiles to RSC depletion known to induce global NDR shrinkage (Kubik et al. 2018). Analyses of published data show a global increase of nucleosome occupancy over the TBS for all the quintiles (Fig. 7F) (Kubik et al. 2018). However, this increase is less important for the genes of the first quintile. Thus, as expected from our model, the genes showing the highest levels of antisense transcription into promoters are less sensitive to RSC depletion because they already present a higher nucleosome occupancy within the NDR at steady-state resulting from decreased interaction of the NDR-flanking nucleosomes with RSC.

Altogether, we propose that the chromatin-driven model of antisense-mediated transcription interference defined with a subset of genes in the inducible system can be extended with high significance up to 20% of the genes. These observations support the view that gene regulation through interleaved noncoding transcription is a major player in global chromatin shaping of promoters and gene expression in *S. cerevisiae*.

## Discussion

### A chromatin-driven transcription interference model derived from an antisense inducible system

Based on our results, we propose an antisense-mediated transcription interference mechanism through chromatin regulation (Fig. 7A). The rescue of the different molecular phenotypes observed at AMRG in the absence of the Rpd3 HDAC enables us to validate such a chromatin-driven model (Fig. 6). However, the rescue is only partial and other parameters have to be taken into consideration.

First antisense-mediated transcription interference may also implicate other HDACs as suggested by different earlier reports indicating the involvement of Set3 through recruitment by H3K4me2 or of Hda1 (Camblong et al. 2007; Kim et al. 2016). Another component of transcriptional interference may be the physical eviction of the sense PIC by the RNAPII travelling in antisense direction. However, we think this is unlikely since many DNA-binding proteins behave as roadblocking factors in front of which RNAPII stalls (Colin et al. 2014; Mayer et al. 2015; Candelli et al. 2018). Nevertheless, the possibility of RNAPII removing DNA-binding factors remains to be thoroughly tested.

The fold-change obtained with the antisense inducible system for some of the molecular phenotypes are of low amplitude, albeit highly significant when comparing AMRG to the Others. Yet, we do not expect drastic changes mainly because AMRG present relatively high levels of natural antisense transcription into promoters and are consequently already lowly expressed at steady state (Supplemental Fig. S6B). Their chromatin is therefore already partially closed before antisense induction and Nrd1 anchor away only results in a subtle additional gain in repressive chromatin or a weak loss of active chromatin (Fig. 1–5; Supplemental Fig. S6B, C). Importantly, if we observe a magnitude of changes from 5-15% at the chromatin level for the AMRG (Fig. 1–5), it ultimately leads to a 40% mRNA decrease as measured by RNA-seq (Fig. 1C). Thus, low amplitude phenotypes at the chromatin level have important consequences on the cellular RNA content.

### Subpopulations of H3K36me3 and H3K18ac nucleosomes are differently positioned

Induction of antisense into AMRG promoters leads to an increase in nucleosome occupancy over the TBS correlating with a loss of occupancy at −1/+1 (Fig. 2). Concomitantly, a subpopulation of *de novo* H3K36me3 nucleosomes appears over the TBS while H3K18ac nucleosomes decrease at −1/+1 (Fig. 3). These considerations are not restricted to AMRG promoters but are applicable to all nucleosomes undergoing forced antisense (Fig. 4). Altogether, these data strongly suggest the existence of subpopulations of H3K36 and H3K18ac nucleosomes that show different positioning.

Based on these observations, we propose a simple model in which the transcription-associated H3K36 methylated nucleosomes prevent transcription initiation while acetylated nucleosomes are pushed aside allowing PIC binding. Thus, gene expression would depend on the metastable state of the promoter oscillating between a closed and open conformation. Antisense transcription frequency may promote NDR closing by modulating the *de novo* H3K36me3 levels at NDR-flanking nucleosomes while histone demethylases, histone exchange or Histone Acetyl Transferases (HATs) may counteract the shrinkage. Similarly, such nucleosome movements are observed upon gene activation during the metabolic cycle when gene expression is synchronized (Nocetti and Whitehouse 2016). In agreement with our model, gene expression is maximal when −1/+1 nucleosomes are pushed aside in a movement mainly driven by H3K9ac and H3K18ac modifications (Weiner et al. 2010; Hughes et al. 2012; Nocetti and Whitehouse 2016; Sanchez-Gaya et al. 2018).

We do not detect a shift of the nucleosome peaks themselves upon antisense induction at AMRG (Fig. 4A), most likely because we are analyzing regions in which transcription occurs on both strands in the population but is dominated in frequency by sense transcription. Thus, nucleosome phasing is mainly dictated by sense transcription and the relative low frequency of antisense production only displaces a subpopulation and not the whole peaks. In other words, antisense induction may increase cell-to-cell variability in nucleosome positioning at AMRG promoters and gene bodies within a chromatin structure mainly imposed by sense transcription.

### Antisense-mediated transcription interference through chromatin regulation as a widespread mechanism of gene expression control in *S. cerevisiae*

Our results indicate that the top 20% of the genes showing the highest levels of antisense transcription into promoters are significantly less expressed than the others when taking into account nascent transcription data (Fig. 7B). However, in agreement with published observations, the global anticorrelation between sense and antisense transcription is low (Supplemental Fig. 6D; Pearson correlation r=-0.13) (Murray et al. 2015; Brown et al. 2018). These analyses indicate that noncoding nascent transcription needs to reach a certain absolute level in the promoter NDR to induce transcription interference as already proposed by (Nevers et al. 2018) using RNA-seq data. When this level is reached, transcription into the promoter NDR becomes a dominant parameter, possibly through the chromatin rearrangement mechanism proposed here. Nevertheless, other parameters not considered in our study are likely to be involved, including different promoter sequences or transcription factors, the presence of roadblocking proteins or the recruitment of chromatin remodelers and histone modifiers.

We show that gene promoters with high levels of natural antisense tend to be more closed as compared to the others, a finding in agreement with published data (Dai and Dai 2012; Murray et al. 2015). However, our results are in contradiction with the observation that high levels of antisense into promoters correlate with low levels of H3K36me3 and high levels of histone acetylation at promoters (Murray et al. 2015; Brown et al. 2018). This difference is mainly due to the normalization procedure as our nucleosome modification data were not normalized to H3 or MNase-seq levels. Considering the shape of the nucleosomal signal along the DNA, normalization in the “valley” region corresponding to the promoter is more sensitive to the background, increasing the probability of a biased result.

The proposed model mainly considers antisense noncoding transcription, however a comparable transcription interference mechanism may take place as a result of upstream *in tandem* noncoding transcription overlapping with a downstream promoter. Indeed, based on the described involvement of both nucleosome positioning and H3K36me3 at specific loci (Martens et al. 2004; Hainer et al. 2011; Kim et al. 2016), our model may be relevant to transcription interference by noncoding transcription in a variety of configurations.

Importantly, our analyzes were only performed in rich medium. Early termination of lncRNA can be regulated in response to growth conditions leading to different patterns of noncoding transcription elongation (Bresson et al. 2017; van Nues et al. 2017). Thus, a different framework of transcription interference may be expected depending on external conditions.

It is worth noting that about a hundred coding genes are less expressed in mRNA orientation than in lncRNA orientation at the nascent transcription level (data not shown). Thus, in some cases, the mRNA may appear as the intragenic transcript of the lncRNA. This concept, even if marginal in proportion, perfectly illustrates the plasticity of transcriptional circuits. RNAPII transcription happens all along the genome and evolution may shape the balance between expression noise and functionality (Struhl 2007).

### A common model for all eukaryotic NDRs?

In *S. cerevisiae*, NDRs are also regions where replication initiates, mainly through the accessibility of the Origin Recognition Complex (ORC) to the ARS (Autonomously Replicating Sequence) Consensus Sequence (ACS) (Lai and Pugh 2017). We recently showed that noncoding transcription entering into an ARS NDR is able to influence replication initiation by closing the NDR through increased nucleosome occupancy, as well as elevated H3K36me3 and decreased H3K18ac (Soudet et al. 2018). Importantly, replication defects induced by noncoding transcription readthrough into ARS NDR can be partially rescued in the absence of Set2. Although we did not have a precise mechanism in our previous study, the similarity between these earlier and the current observations tend to converge to a unique model that may be applicable to all types of NDRs.

Such a general mechanism may also be relevant to higher eukaryotes. Interestingly, in mammalian cells, replication mainly initiates within gene promoters (Miotto et al. 2016; Chen et al. 2019). Replication initiation efficiency nicely correlates with transcription initiation strength, which itself correlates with the accessibility of the NDR (Brown et al. 2018; Chen et al. 2019). Similarly to yeast cells, high levels of antisense lncRNAs over promoter NDRs correlate with a transcription interference phenotype and a closed NDR conformation (Chen et al. 2016; Brown et al. 2018). Together, these observations suggest that noncoding transcription readthrough into NDRs may be a neglected parameter regulating both replication initiation and gene expression in mammalian cells through a chromatin-driven mechanism. It would therefore also be of interest to analyze the consequences of noncoding transcription readthrough on the chromatin structure of enhancers which also correspond to NDRs. High resolution maps of nucleosomes and histone modifications will reveal whether the proposed chromatin-driven model can be extended to mammalian cells.

## Acknowledgements

We thank Guillaume Canal, Charlie Rochat, Florian Steiner, Michel Strubin and all members of the Stutz laboratory for critical reading of the manuscript, comments and discussions. We thank Mylene Docquier and the iGE3 genomics platform of the University of Geneva for performing all the deep sequencing. This work was supported by funds from the Swiss National Science Foundation (grants 31003A_153331 and 31003A_182344 to F.S.), iGE3 and the Canton of Geneva.

## Author Contributions

J.K.G, A.M., V.G.M. and J.S. performed the experiments, J.KG, F.S. and J.S. analyzed the data and F.S. and J.S. conceived and supervised the study. J.K.G, F.S. and J.S. interpreted the data. J.S. wrote the manuscript with input from all authors.

## Methods

### Saccharomyces cerevisiae strains and growth

All strains were derived from the Anchor-Away genetic backgrounds (see Supplemental Table 1) (Haruki et al. 2008). Cells were grown in YEPD medium (1% yeast extract, 1% peptone) supplemented with 2% glucose as carbon source. All strains were grown at 30°C and were not affected in growth by the different tags or deletions (Supplemental Table 1). Anchor-away of Nrd1-AA was induced by adding 1µg/ml of rapamycin to the medium.

### RNA extraction and RNA-seq

RNAs were extracted using Glass-beads and TRIzol (Invitrogen). RNA library preparation and single-end stranded sequencing were performed at the IGE3 genomics platform of the University of Geneva.

### MNase(-ChIP)-seq

The MNase- and MNase-ChIP-seq experiments were performed mainly as described in (Weiner et al. 2015) with the following modifications. Nrd1-AA strain was inoculated overnight and diluted in the morning to OD_600_=0.2 in YEPD medium. Yeast cell cultures of 100 ml was used per modification. Rapamycin treatment was performed at OD_600_=0.5 for 60 minutes on half of the culture. The cells were fixed with formaldehyde at a final concentration of 1% for 15 minutes followed by glycine addition at a final concentration of 125 mM for 5 minutes. Cells were washed 2x with ddH_2_O, harvested in magnaLyser tubes (Roche) (100 mL cell culture/tube) and frozen at −20°C.

Chromatin extraction was performed by breaking cells in 1 mL of Cell Breaking buffer (0.1 M Tris-HCl, pH 7.9, 20% glycerol, EDTA-free protease inhibitors) and 1 mL of acid-washed glass beads in a magnaLyser (Roche). The cell extract was then centrifuged at 19,000g at 4 °C for 10 minutes to collect chromatin. The chromatin was resuspended in 600 μL of NP buffer (0.5 mM Spermidine, 50 mM NaCl, 1mM β-mercaptoethanol, 0.075% NP-40, 10 mM Tris-HCl pH 7.4, 5 mM MgCl_2_, 1 mM CaCl_2_) per tube. The DNA concentration was measured using Qubit dsDNA BR assay kit (Invitrogen). Chromatin was diluted to 20 μg/mL using NP buffer in all conditions for comparable results.

MNase treatment was performed using 0.2 μL of MNase (ThermoScientific 100 units/μL) / 12 μg of chromatin in 600 μL reaction mixture for 14 minutes at 37 °C. To perform all the modifications for one replicate from one cell culture the reaction was scaled 10x. The reaction was stopped by adding EDTA (final concentration 20 mM) in the reaction mixture.

For MNase-seq, an equal amount of elution buffer (10 mM Tris-HCl, pH 8.0, 1% SDS, 150 mM NaCl, 5 mM DTT) was added. The further steps for MNase-seq are the same as those performed after MNase-ChIP elution.

For MNase-ChIP, the buffer of the MNase reaction mixture was adjusted to be compatible to FA-lysis buffer (50 mM HEPES-KOH, pH 7.5, 140 mM NaCl, 1 mM EDTA, 1% Triton X-100, 0.1% sodium deoxycholate, EDTA-free protease tablet). The following salts were added to the MNase reaction tube: 80 μL of 0.5 M HEPES-KOH, pH 7.5, 22.4 μL of 5 M NaCl pre-mixed with protease inhibitors. This was followed by the addition of detergents, 6.4 μL of 12.5% sodium deoxycholate and 80 μL of 10% Triton X-100 per 600 μL MNase reaction.

The protein-G dynabeads (ThermoFisher Scientific) were pre-incubated with the antibodies for one hour at 4°C on a rotor, 85 μL beads/reaction. The antibodies used were anti-H3 (Abcam ab1791):12 μL per reaction; anti-H3K36me3 (Abcam ab9050): 6 μL per reaction; anti-H3K18ac (abcam ab1191): 7 μL per reaction; H4ac (Merck Millipore 06-598): 10 μL per reaction. The beads were washed in FA-lysis buffer twice, followed by incubation with the MNase-treated chromatin for 4 hours at 4°C on a rotor. The beads were sequentially washed with 1 mL of FA-Lysis buffer, 1 mL FA500-lysis Buffer (FA-Lysis Buffer + 500 mM NaCl), 1 mL of Buffer III (10 mM Tris-HCl, pH 8.0, 1 mM EDTA, 250 mM LiCl, 0.5% NP-40, 0.05% sodium deoxycholate), 1 mL of Tris-EDTA pH 8.0. For elution, the beads were incubated in 150 μL of elution buffer at 65 °C for 15 minutes.

The crosslinking was reversed by incubation of the eluate at 65°C overnight followed by the addition of 150 μL of TE (10 mM Tris-HCl, pH 8.0, 1 mM EDTA). The eluate was then treated with proteinase K (1 μg/μL final concentration) for 3 hours at 42°C. The DNA was purified using the NucleoSpin Gel and PCR clean-up kit (Macherey-Nagel) using the SDS protocol provided in the manual. The DNA concentration was measured using Qubit dsDNA HS assay kit (Invitrogen) and the libraries were prepared using NEBnext Ultra DNA library prep kit for Illumina (NEB). Finally, samples were paired-end sequenced at the iGE3 genomics platform of the University of Geneva.

### ChEC-seq

The experiment was performed as described in (Zentner et al. 2015) with some modifications. The yeast strains were cultured the day before in YEPD medium. They were diluted to OD_600_=0.2 in YEPD medium in the morning. The 60 min rapamycin treatment was performed at OD_600_=0.4.

For each condition, 50 mL cell cultures were harvested at room temperature. The cells were washed twice in 1 mL Buffer A (15 mM Tris-HCl pH 7.5, 80 mM KCl, 0.1 mM EGTA, 0.2 mM spermine, 0.5 mM spermidine and protease inhibitor tablet (Sigma-Aldrich)). Cells were resuspended in 594 μL of buffer A and 6 μL of 10% digitonin (0.1% final concentration) were added to permeabilize the cells during 5min at 30°C. CaCl_2_ was added (final concentration 5 mM) to activate the MNase cleavage. 200 μL were collected at 30s for *SPT15-MNase* or 20s for *STH1-MNase* and were immediately mixed with 200 μL of 2X Stop solution (400 mM NaCl, 20 mM EDTA, 4 mM EGTA, 1% SDS).

The cells were treated with Proteinase K (0.4 μg/μL final concentration) and incubated at 55°C for 30 minutes. The DNA was extracted using phenol:chloroform:isoamyl extraction, and precipitated by adding 30 μg glycogen, 500 μL of 100% ethanol and incubated at −20°C for one hour. RNAse treatment was performed by adding 34.5 μL of Tris, pH 8.0 and 10 μg of RNAse per reaction.

For size selection of DNA fragments, 75 μL of solid phase reversible immobilization beads (SPRI, AmpureXP Beckman Coulter) were added in 25 μL of RNAse treated DNA and the reaction was mixed by pipetting up and down 10 times. The beads were incubated at room temperature for 5 minutes. The tubes were placed in the magnetic rack and supernatant was transferred to the new tube containing 10 mM Tris and 0.2 M NaCl for each reaction. The DNA was extracted with phenol:chloroform:isoamyl solution and precipitated with 100% ethanol. The pellets were washed with 70% ethanol and resuspended in 29 μL of Tris, pH 8.0. DNA concentration was measured using Qubit dsDNA HS assay kit (Invitrogen) using 4 μL sample. The remaining 25 μL were used to prepare sequencing libraries using NEBnext Ultra DNA library prep kit for Illumina (NEB). The samples were sequenced in the paired-end mode at the iGE3 genomics sequencing platform in Geneva.

### ATAC-seq

The experiment was performed after modifying the protocol from (Schep et al. 2015). The yeast strains were inoculated the day before in YEPD. Cultures were diluted to OD_600_=0.2 and allowed to reach OD_600_=0.4. Half of the culture was then treated with Rapamycin for 1h at 30°C.

1 mL of the cell culture was harvested from all conditions and strains. Cells were washed twice in 1 mL sorbitol buffer (1.4 M Sorbitol, 40 mM HEPES-KOH, pH 7.5, 0.5 mM MgCl_2_), resuspended in 190 μL of sorbitol buffer and 10 μL of 10 mg/mL Zymolyase (20T Zymolyase, AMSBIO), and incubated at 30°C shaking at 200 rpm for 30 minutes. The cells were washed twice in sorbitol buffer and resuspended in 95 μL of 1x TD buffer and 5 μL of the TD enzyme (Illumina). The reaction was incubated at 37 °C at 500 rpm for 30 minutes. DNA was then extracted with 3x SPRI beads (AmpureXP Beckman Coulter) and eluted in 28 μL 10 mM Tris-HCl, pH 8.0.

Initial PCR amplification was performed with 4 μL of transposed DNA to check the transpositions and cycles required. Final amplification was performed with NEBNext High-Fidelity 2x PCR Master Mix (NEB) using Nextera Primers (Illumina). 12 cycles were performed.

The size selection was performed by sequential SPRI beads precipitation. First, the DNA was mixed by pipetting in 0.5x beads followed by incubation for 5 minutes at room temperature. The supernatant was used for the next DNA precipitation with 2.5x SPRI beads. The supernatant was discarded and the beads were washed with 70% ethanol twice before the DNA was extracted using 28 μL of 10 mM Tris-HCl, pH 8.0. The DNA concentration was measured and the samples were paired-end sequenced at the iGE3 genomics platform of the University of Geneva.

### ChIP-qPCR

The experiments were performed as described in (Kuras and Struhl 1999) after some modifications. For both *S. cerevisiae* and *S. pombe* (used here for spike-in), the cultures were grown to exponential phase until OD_600_=0.5, after which they were treated with Rapamycin for different time points. Cells were fixed and chromatin was extracted as in the MNase(-ChIP)-seq section with the exception of the breaking step performed in FA-lysis buffer (50 mM HEPES-KOH, pH 7.5, 140 mM NaCl, 1 mM EDTA, 1% Triton X-100, 0.1% sodium deoxycholate, EDTA-free protease tablet). Chromatin was sheared to 300 bp fragments through sonication (Bioruptor, Diagenode).

2/3^rd^ *S. cerevisiae* chromatin was mixed with 1/3^rd^ of *S. pombe* chromatin. The ChIP was performed with the mixture using on average 200 μL of the ChIP chromatin at a concentration of 20 μg/mL. The rest of the experiment was performed as in the MNase(-ChIP)-seq section. After elution and clean-up, DNA fragments were amplified with the different oligos of Suppl. Table 2 using the SYBR Green PCR Master Mix (Applied Biosystems) and a Real-Time PCR machine (Bio-Rad).

### List of genes, TBS and nucleosomes

The list of gene coordinates from TSS to poly-A was kindly provided by the Mellor Lab. Among them were picked the ones considered as “Verified” genes in the Saccharomyces Genome Database (SGD) giving a complete list of 4,775 coding genes. For the TBS coordinates, our list was crossed with the data from the Pugh lab (Rhee and Pugh 2012). −1/+1 coordinates were extracted from DANPOS2 analysis with default settings (Chen et al. 2013) from our Wig profile of H3K18ac in -Rap. Figure 4A was obtained using the coordinates of the nucleosome atlas from the Friedman lab (Weiner et al. 2015).

### RNA-seq analysis

Single-end reads were aligned to sacCer3 genome assembly using Bowtie2 (Langmead and Salzberg 2012) with options ‘-k 20 --end-to-end --sensitive -X 800’. PCR duplicates were removed from the analysis. BigWig coverage files were generated using Bam2wig function. Differential expression analysis was performed using the R/Bioconductor package DEseq on mRNA annotations Ensembl (Saccharomyces_cerevisiae.EF4.65.gtf). Antisense transcripts with a fold-change of at least 2 and multiple testing adjusted p-value lower than 0.05 were considered differentially expressed. Among them, AMRG were defined as the genes showing a <0.8 fold-change with an adjusted p-value < 0.05.

### MNase(-ChIP)-seq mapping

Paired-end reads were aligned to sacCer3 genome assembly using Bowtie2 (Langmead and Salzberg 2012) with options ‘-k 20 --end-to-end --sensitive -X 800’. PCR duplicates were removed from the analysis. Then, deepTools 2.0 (Ramirez et al. 2016) was used through the bamCoverage function with size selection of fragments (120-200bp to visualize only proper nucleosomes and not “fragile nucleosomes” (Kubik et al. 2015; Brahma and Henikoff 2019)), counting only the 3bp at the center of fragments and cpm normalization.

### ChEC-seq and ATAC-seq mapping

Adapters were first removed from the paired-end reads using the TrimGalore! Tool with default options from the Galaxy server (Afgan et al. 2018). Paired-end reads were then aligned to sacCer3 genome assembly using Bowtie2 (Langmead and Salzberg 2012). PCR duplicates were removed from the analysis. DeepTools 2.0 (Ramirez et al. 2016) was then used through the bamCoverage function with size selection of fragments (0-120bp for TBP-ChEC and ATAC-seq, and 120-200bp for Sth1-ChEC) and counting of only the 3bp at the center of fragments.

### Metagene analyses

Bigwig files of independent duplicates generated *via* mapping were then averaged in deepTools2.0 using the bigWigCompare command (however, results of each duplicate are shown in Supplemental Fig.). Metagene plots were produced using computeMatrix followed by plotProfile commands. +Rap/-Rap fold-changes over TBS or −1/+1 nucleosomes were calculated adding no pseudo-count for ATAC-seq, TBP-ChEC and Sth1-ChEC, a pseudo-count of 1 for RNAPII PAR CLIP and a pseudo-count of 0.01 for MNase-seq, H3K36me3, H3K18ac and H4ac marks.

Fuzziness was calculated with DANPOS2 (Chen et al. 2013) using H3K18ac -Rap and H3K36me3 -Rap Wig files. Only peaks showing >15cpm were taken into account in the analysis (7’929 and 9’899 peaks for H3K36me3 and H3K18ac, respectively).

Fig. 6A-D were normalized as follows: the mean of +Rap/-Rap fold-change was normalized to 1 in both *WT* and *rpd3Δ* strains giving a normalization factor for each strain. These normalization factors were then applied to the different classes to correct the raw fold-changes.

### Statistical analyses

All plots and statistical analyses of this work were performed using Prism 8.0 (Graphpad). All tests are nonpaired tests (with the exception of Fig. 6A-D based on paired tests). *t*-tests or Mann–Whitney *U* tests were used according to the normality of the data analyzed, which was calculated using a d’Agostino-Pearson omnibus normality test. * if p-value < 0.05, **< 0.01, *** < 0.001, ****<0.0001.

### Data availability

The accession number for the data reported in this study is GEO: GSE130946. RNAPII PAR CLIP data in Nrd1-AA were retrieved from (Schaughency et al. 2014). Nucleosome profiles in the TBP-AA and Sth1-AA strain were generated by (Kubik et al. 2018) and (Tramantano et al. 2016).

---

**Supplemental Figure S1: related to Figure 1.**
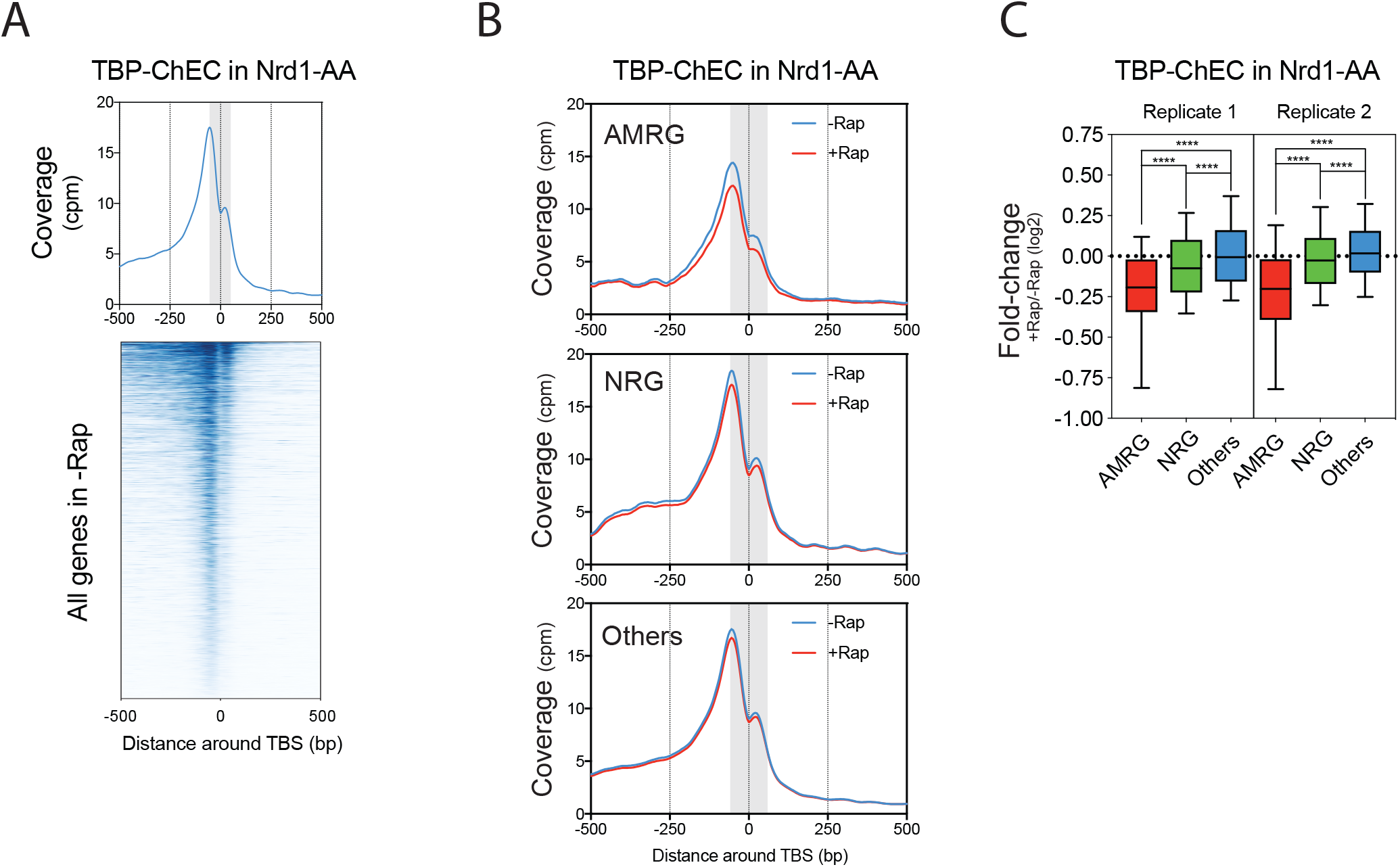
**(A)** Metagene analysis and heatmap depicting the TBP-ChEC profile in the Nrd1-AA in the absence of Rap. Results are centered on the TATA-Binding Site (TBS). **(B)** Metagene analyses of TBP-ChEC induced for 30sec in an Nrd1-AA strain treated or not with Rap for 1h. The grey box represents the 50bp area around the TBS over which statistics are generated. The center of 0-120bp paired-end fragments is represented for the plots. **(C)** Box-plots of the two independent replicates of TBP-ChEC fold-change related to Figure 1G. The fold-change is measured over a 50bp region centered on the TBS.

**Supplemental Figure S2: related to Figure 2.**
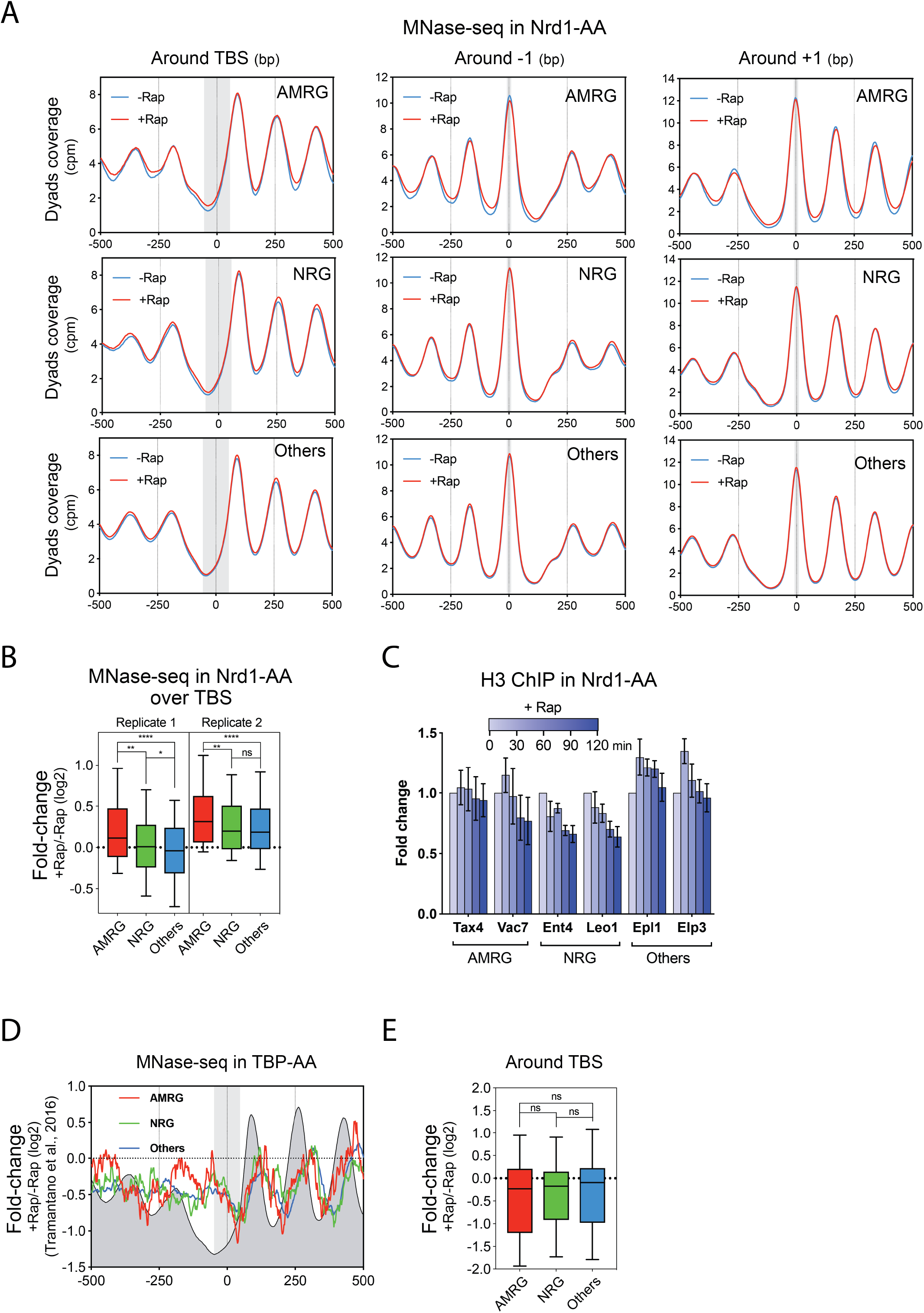
**(A)** Aggregate plots centered on TBSs, −1 and +1 nucleosomes, respectively, of MNase-seq profiles obtained in an Nrd1-AA strain treated or not for 1h with Rap. The centers of 120-200bp paired-end fragments are represented. Grey rectangles represent the 50bp-TBS centered and 10bp--1/+1 nucleosomes-centered areas. **(B)** Box-plots of the two independent replicates of MNase-seq fold-change related to Figure 2B. The fold-change is measured over a 50bp region centered on the TBS. **(C)** ChIP of H3 at gene promoters of AMRG, NRG and Other genes. ChIPs were performed at 0, 30, 60, 90 and 120min after rapamycin addition. Immunoprecipitated promoter NDRs were normalized to immunoprecipitated *SPT1*5 ORF after qPCR amplification. Primers are designed to target promoter NDRs. The fold-change was artificially set to 1 for each gene in the -Rap condition. Error bars represent the Standard Error of the Mean (SEM) for a set of 3 independent experiments. **(D)** Metagene plot of MNase-seq performed in a TBP-AA strain treated or not with Rap for 30min. Midpoint of 120-200bp are represented. The grey box represents the 50bp area centered on the TBS. Results were retrieved from (Tramantano et al., 2016). **(E)** Box-plots of the dyads occupancy fold-change in +Rap/-Rap in the TBP-AA strain. The fold-change is measured over a 50bp region centered on the TBS.

**Supplemental Figure S3: related to Figure 3.**
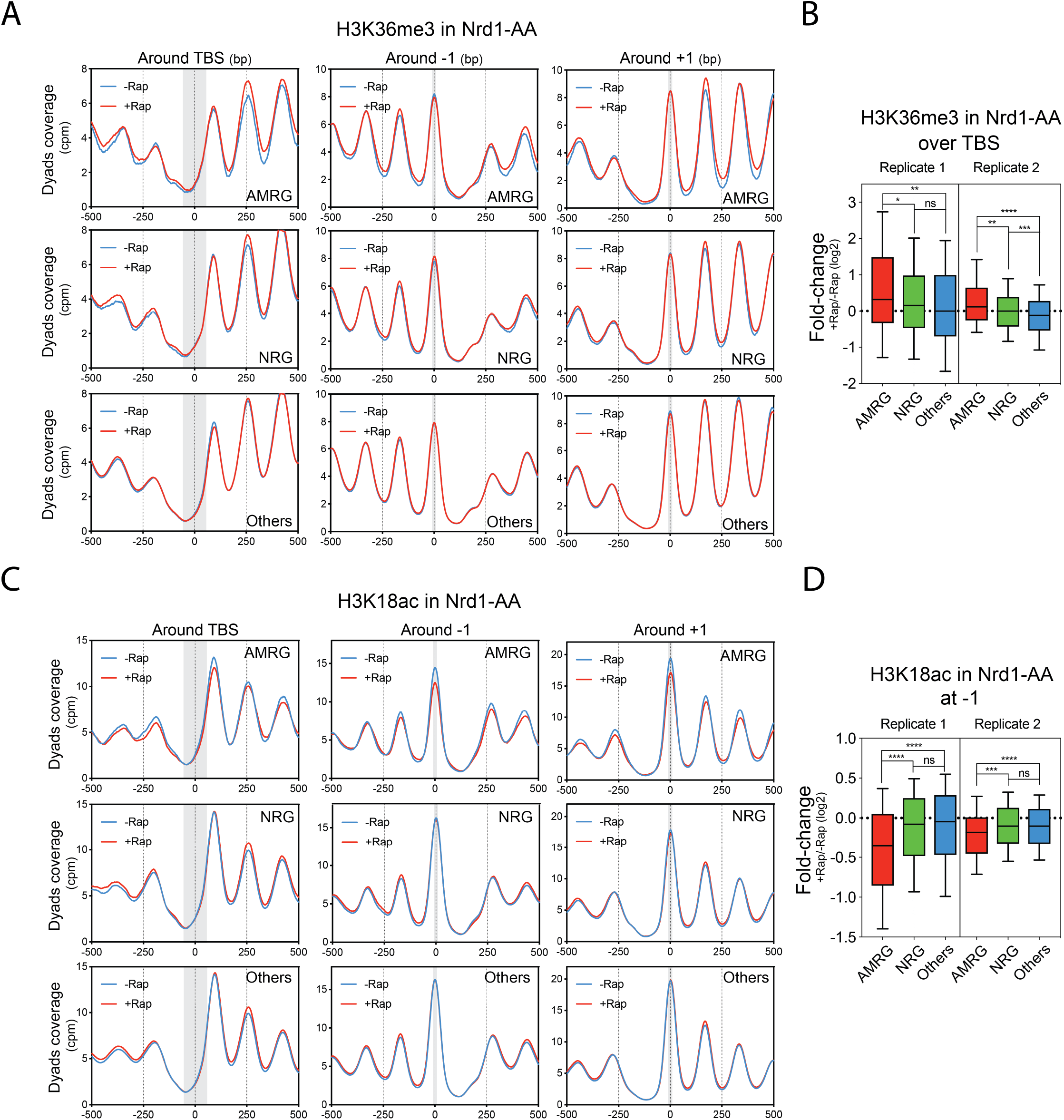
**(A)** Metagene plot of H3K36me3 MNase-ChIP-seq related to Figure 3A and centered on the TBS, −1 and +1 nucleosomes, respectively. Midpoint of 120-200bp fragments are represented. **(B)** Box-plots of the fold-change related to Figure 3B for the two independent replicates of H3K36me3 MNase-ChIP-seq. The fold-change is measured over a 50bp region centered on the TBS. **(C)** Metagene plot of H3K18ac MNase-ChIP-seq related to Figure 3C and centered on the TBS, −1 and +1 nucleosomes, respectively. Midpoint of 120-200bp fragments are represented. **(D)** Box-plots of the fold-change related to Figure 3D for the two independent replicates of H3K18ac MNase-ChIP-seq. The fold-change is measured over a 10bp region centered on the −1 nucleosome.

**Supplemental Figure S4: related to Figure 5.**
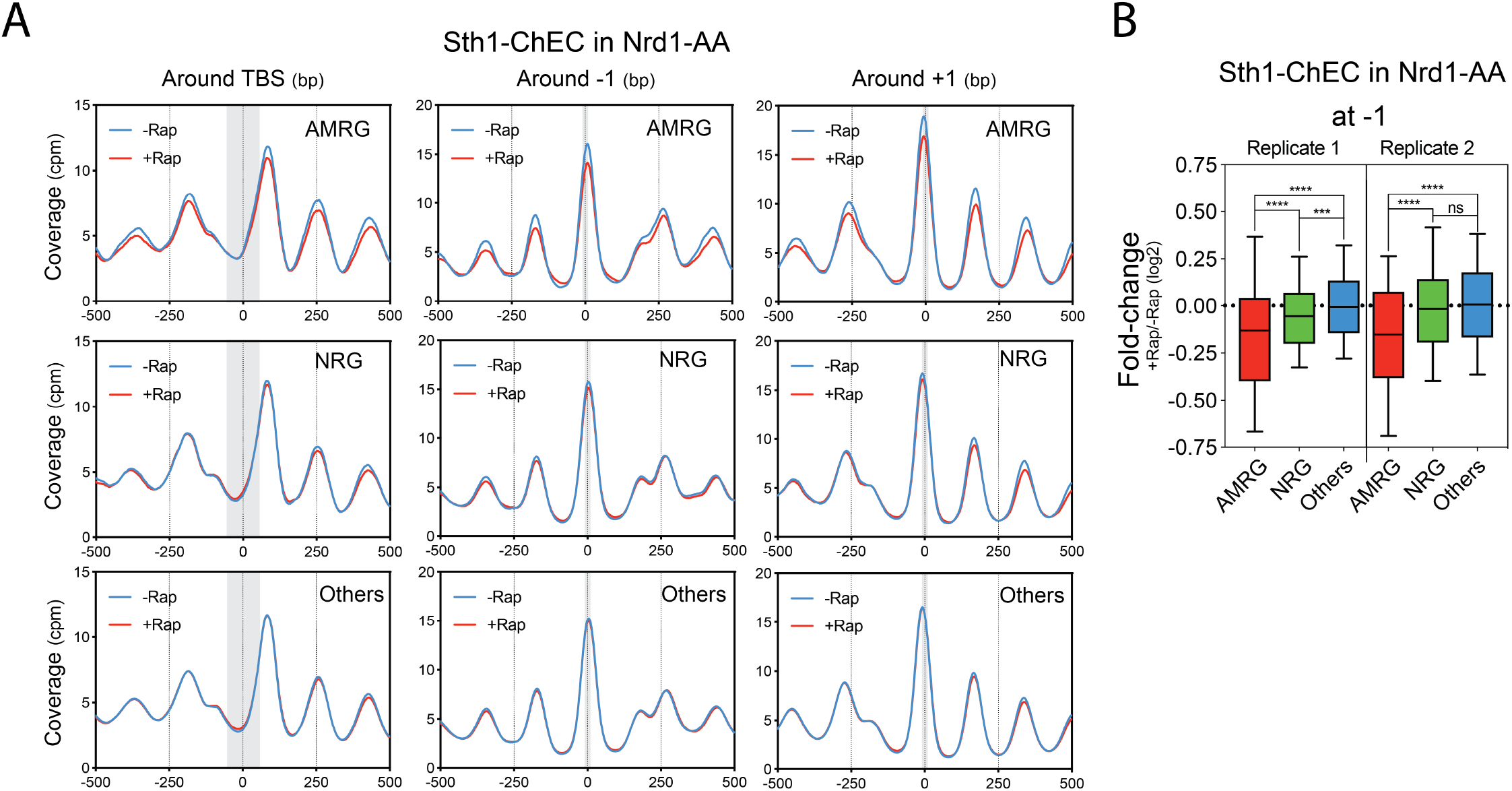
**(A)** Metagene plot of Sth1-ChEC centered on the TBS, −1 and +1 nucleosomes, respectively. Midpoints of 120-200bp fragments are represented. **(B)** Box-plots of the (+Rap/-Rap) fold-change related to Figure 5B for the two independent replicates of Sth1-ChEC. The fold-change is measured over a 10bp region centered on the −1 nucleosome.

**Supplemental Figure S5: related to Figure 6.**
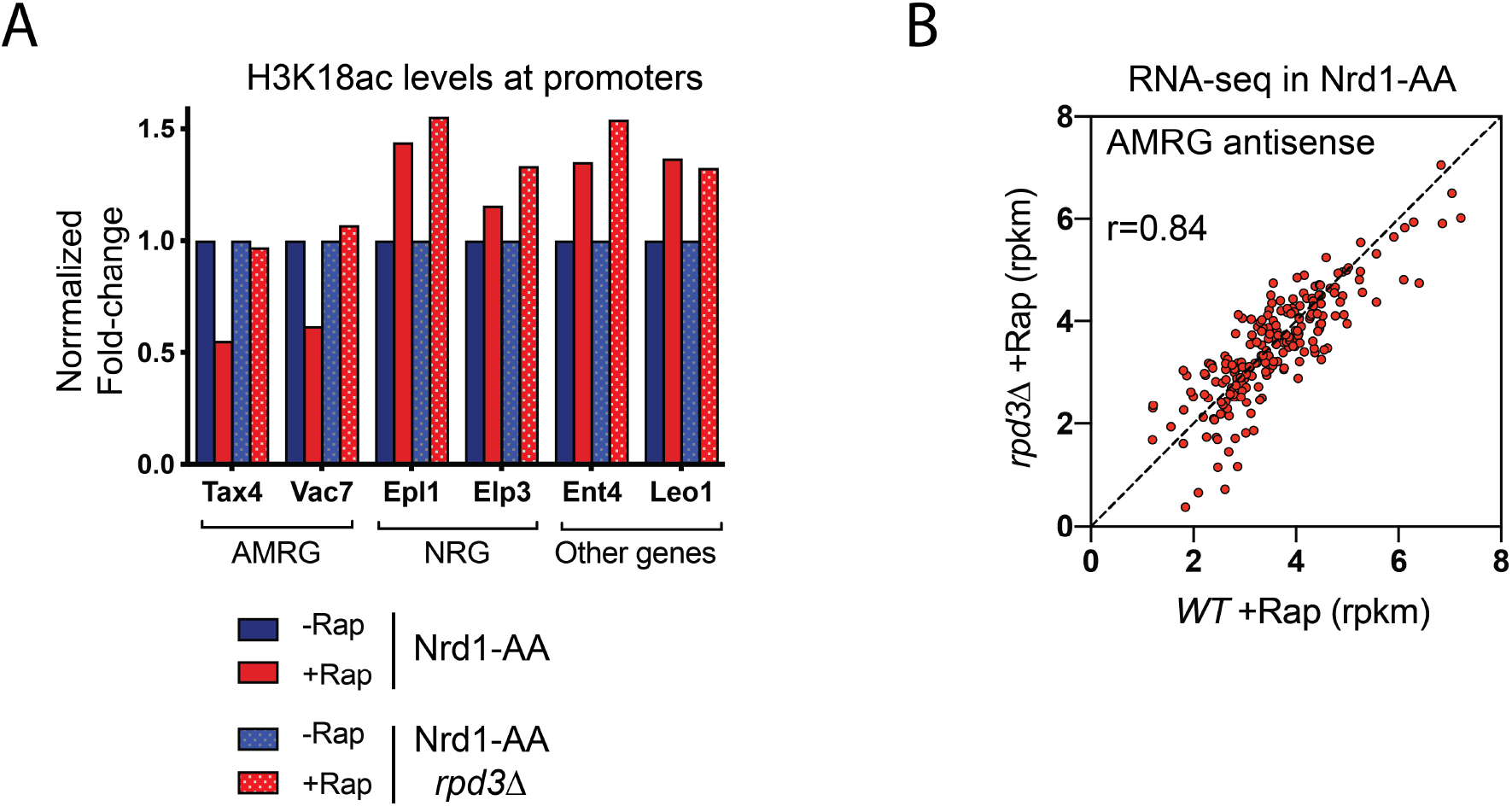
**(A)** ChIP of H3K18ac levels at the promoters of AMRG, NRG and Other genes. *S. pombe* chromatin was used as a spike-in control and mixed with *S. cerevisiae* chromatin before immunoprecipitation. *S. cerevisiae* results are normalized to *S. pombe ACT1* ORF. All results are expressed as fold-change with respect to -Rap which value was set to 1. **(B)** RNA-seq of AMRG antisense in Nrd1-AA +Rap vs Nrd1-AA *rpd3Δ* +Rap. Results are depicted in rpkm. The +Rap condition was chosen to have an accurate measurement of antisense production. r indicates the Pearson correlation. Even in the absence of Rpd3, antisense RNAs are globally well produced at AMRGs and the rescue of AMRG sense expression (Figure 6) is not a consequence of a lack of antisense production.

**Supplemental Figure 6: related to Figure 7.**
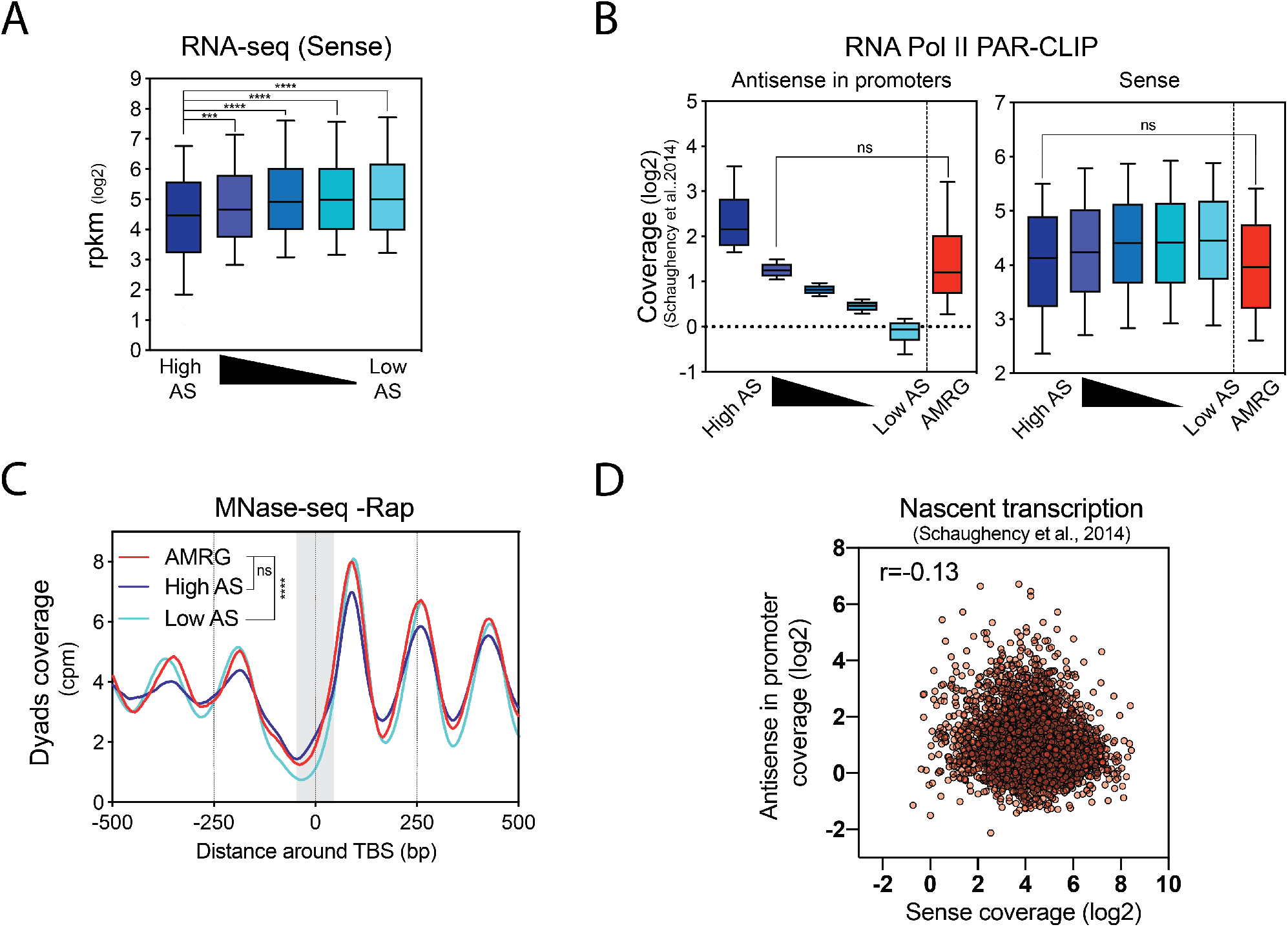
**(A)** Box-plots of sense expression by RNA-seq according to the quintiles defined in Figure 7. **(B)** Natural nascent antisense levels in promoters and natural nascent sense levels of AMRG as compared to the quintiles defined in Figure 7. **(C)** Metagene plot of MNase-seq profile at steady state for the AMRG as compared with the high and low antisense quintiles defined in Figure 7. Results are centered on the TBS. **(D)** Global correlation between nascent antisense transcription into promoters (−100bp to TSS area) and nascent sense transcription (100bp area upstream of the poly-A site). The correlation coefficient r corresponds to the Pearson correlation.

**Suppl. Table 1.**

| Experimental Models:<br>Organisms/Strains |  |  |
| --- | --- | --- |
| *S. cerevisiae* AA<br>(FSY4885) | (Haruki et al.<br>2008) | *MAT α, tor1-1, fpr1Δ::NAT, RPL13A-2×FKBP12::TRP1* |
| *S. cerevisiae* Nrd1-AA<br>(FSY5065) | (Castelnuovo<br>et al. 2014) | *MAT α, tor1-1, fpr1Δ::NAT, RPL13A-2×FKBP12::TRP1, NRD1-FRB::KanMX6* |
| *S. cerevisiae* Nrd1-AA<br>*rpd3Δ* (FSY7015) | This study | *MAT α, tor1-1, fpr1::NAT, RPL13A-2×FKBP12::TRP1, NRD1-FRB::KanMX6, rpd3Δ::HIS3* |
| *S. cerevisiae* Nrd1-AA<br>TBP-MNase (FSY8162) | This study | *MAT α, tor1-1, fpr1::NAT, RPL13A-2×FKBP12::TRP1, NRD1-FRB::KanMX6, TBP-MNase::HPHMX6* |
| *S. cerevisiae* Nrd1-AA<br>*rpd3Δ* TBP-MNase<br>(FSY8164) | This study | *MAT α, tor1-1, fpr1::NAT, RPL13A-2×FKBP12::TRP1, NRD1-FRB::KanMX6, rpd3Δ::HIS3, TBP-MNase::HPHMX6* |
| *S. cerevisiae* Nrd1-AA<br>STH1-MNase<br>(FSY8254) | This study | *MAT α, tor1-1, fpr1::NAT, RPL13A-2×FKBP12::TRP1, NRD1-FRB::KanMX6, STH1-MNase::HPHMX6* |
| *S. cerevisiae* Nrd1-AA<br>*rpd3Δ* STH1-MNase<br>(FSY8255) | This study | *MAT α, tor1-1, fpr1::NAT, RPL13A -2×FKBP12::TRP1, NRD1 -FRB::KanMX6, rpd3Δ::HIS3, STH1 -MNase::HPHMX6* |

**Suppl. Table 2.**
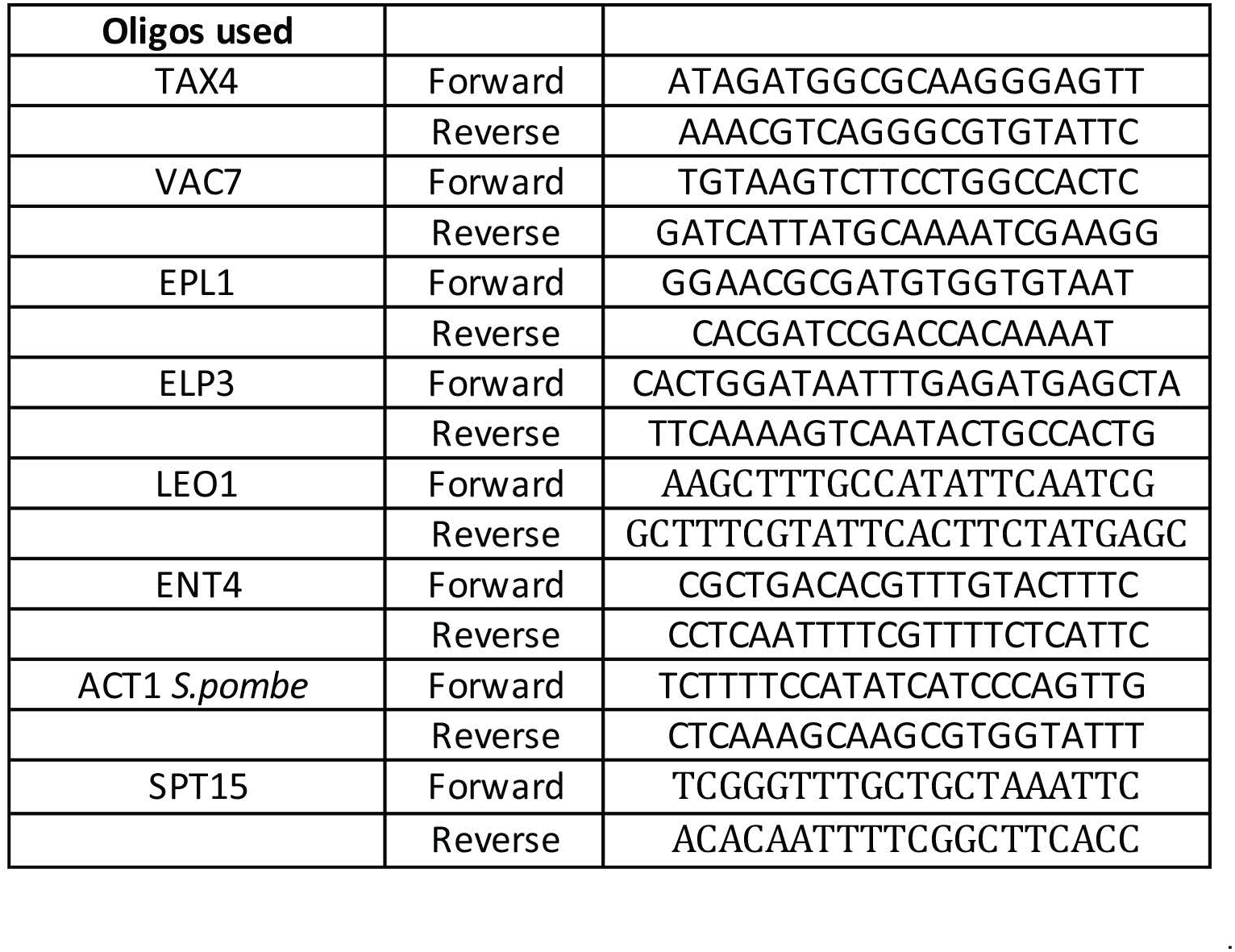

## References

Afgan E, Baker D, Batut B, van den Beek M, Bouvier D, Cech M, Chilton J, Clements D, Coraor N, Gruning BA et al. 2018. The Galaxy platform for accessible, reproducible and collaborative biomedical analyses: 2018 update. Nucleic Acids Res 46: W537–W544.

Badis G, Chan ET, van Bakel H, Pena-Castillo L, Tillo D, Tsui K, Carlson CD, Gossett AJ, Hasinoff MJ, Warren CL et al. 2008. A library of yeast transcription factor motifs reveals a widespread function for Rsc3 in targeting nucleosome exclusion at promoters. Mol Cell 32: 878–887.

Brahma S, Henikoff S. 2019. RSC-Associated Subnucleosomes Define MNase-Sensitive Promoters in Yeast. Mol Cell 73: 238–249 e233.

Bresson S, Tuck A, Staneva D, Tollervey D. 2017. Nuclear RNA Decay Pathways Aid Rapid Remodeling of Gene Expression in Yeast. Mol Cell 65: 787–800 e785.

Brown T, Howe FS, Murray SC, Wouters M, Lorenz P, Seward E, Rata S, Angel A, Mellor J. 2018. Antisense transcription-dependent chromatin signature modulates sense transcript dynamics. Mol Syst Biol 14: e8007.

Camblong J, Iglesias N, Fickentscher C, Dieppois G, Stutz F. 2007. Antisense RNA stabilization induces transcriptional gene silencing via histone deacetylation in S. cerevisiae. Cell 131: 706–717.

Candelli T, Challal D, Briand JB, Boulay J, Porrua O, Colin J, Libri D. 2018. High-resolution transcription maps reveal the widespread impact of roadblock termination in yeast. EMBO J 37.

Carrozza MJ, Li B, Florens L, Suganuma T, Swanson SK, Lee KK, Shia WJ, Anderson S, Yates J, Washburn MP et al. 2005. Histone H3 methylation by Set2 directs deacetylation of coding regions by Rpd3S to suppress spurious intragenic transcription. Cell 123: 581–592.

Castelnuovo M, Rahman S, Guffanti E, Infantino V, Stutz F, Zenklusen D. 2013. Bimodal expression of PHO84 is modulated by early termination of antisense transcription. Nat Struct Mol Biol 20: 851–858.

Castelnuovo M, Zaugg JB, Guffanti E, Maffioletti A, Camblong J, Xu Z, Clauder-Munster S, Steinmetz LM, Luscombe NM, Stutz F. 2014. Role of histone modifications and early termination in pervasive transcription and antisense-mediated gene silencing in yeast. Nucleic Acids Res 42: 4348–4362.

Chatterjee N, Sinha D, Lemma-Dechassa M, Tan S, Shogren-Knaak MA, Bartholomew B. 2011. Histone H3 tail acetylation modulates ATP-dependent remodeling through multiple mechanisms. Nucleic Acids Res 39: 8378–8391.

Chen K, Xi Y, Pan X, Li Z, Kaestner K, Tyler J, Dent S, He X, Li W. 2013. DANPOS: dynamic analysis of nucleosome position and occupancy by sequencing. Genome Res 23: 341–351.

Chen Y, Pai AA, Herudek J, Lubas M, Meola N, Jarvelin AI, Andersson R, Pelechano V, Steinmetz LM, Jensen TH et al. 2016. Principles for RNA metabolism and alternative transcription initiation within closely spaced promoters. Nat Genet 48: 984–994.

Chen YH, Keegan S, Kahli M, Tonzi P, Fenyo D, Huang TT, Smith DJ. 2019. Transcription shapes DNA replication initiation and termination in human cells. Nat Struct Mol Biol 26: 67–77.

Chereji RV, Ramachandran S, Bryson TD, Henikoff S. 2018. Precise genome-wide mapping of single nucleosomes and linkers in vivo. Genome Biol 19: 19.

Churchman LS, Weissman JS. 2011. Nascent transcript sequencing visualizes transcription at nucleotide resolution. Nature 469: 368–373.

Colin J, Candelli T, Porrua O, Boulay J, Zhu C, Lacroute F, Steinmetz LM, Libri D. 2014. Roadblock termination by reb1p restricts cryptic and readthrough transcription. Mol Cell 56: 667–680.

Core LJ, Waterfall JJ, Lis JT. 2008. Nascent RNA sequencing reveals widespread pausing and divergent initiation at human promoters. Science 322: 1845–1848.

Dai Z, Dai X. 2012. Antisense transcription is coupled to nucleosome occupancy in sense promoters. Bioinformatics 28: 2719–2723.

Djebali S, Davis CA, Merkel A, Dobin A, Lassmann T, Mortazavi A, Tanzer A, Lagarde J, Lin W, Schlesinger F et al. 2012. Landscape of transcription in human cells. Nature 489: 101–108.

du Mee DJM, Ivanov M, Parker JP, Buratowski S, Marquardt S. 2018. Efficient termination of nuclear lncRNA transcription promotes mitochondrial genome maintenance. Elife 7.

Hainer SJ, Pruneski JA, Mitchell RD, Monteverde RM, Martens JA. 2011. Intergenic transcription causes repression by directing nucleosome assembly. Genes Dev 25: 29–40.

Hartley PD, Madhani HD. 2009. Mechanisms that specify promoter nucleosome location and identity. Cell 137: 445–458.

Haruki H, Nishikawa J, Laemmli UK. 2008. The anchor-away technique: rapid, conditional establishment of yeast mutant phenotypes. Mol Cell 31: 925–932.

Hughes AL, Jin Y, Rando OJ, Struhl K. 2012. A functional evolutionary approach to identify determinants of nucleosome positioning: a unifying model for establishing the genome-wide pattern. Mol Cell 48: 5–15.

Jensen TH, Jacquier A, Libri D. 2013. Dealing with pervasive transcription. Mol Cell 52: 473–484.

Joshi AA, Struhl K. 2005. Eaf3 chromodomain interaction with methylated H3-K36 links histone deacetylation to Pol II elongation. Mol Cell 20: 971–978.

Kaikkonen MU, Adelman K. 2018. Emerging Roles of Non-Coding RNA Transcription. Trends Biochem Sci 43: 654–667.

Kasten M, Szerlong H, Erdjument-Bromage H, Tempst P, Werner M, Cairns BR. 2004. Tandem bromodomains in the chromatin remodeler RSC recognize acetylated histone H3 Lys14. EMBO J 23: 1348–1359.

Keogh MC, Kurdistani SK, Morris SA, Ahn SH, Podolny V, Collins SR, Schuldiner M, Chin K, Punna T, Thompson NJ et al. 2005. Cotranscriptional set2 methylation of histone H3 lysine 36 recruits a repressive Rpd3 complex. Cell 123: 593–605.

Kim JH, Lee BB, Oh YM, Zhu C, Steinmetz LM, Lee Y, Kim WK, Lee SB, Buratowski S, Kim T. 2016. Modulation of mRNA and lncRNA expression dynamics by the Set2-Rpd3S pathway. Nat Commun 7: 13534.

Klein-Brill A, Joseph-Strauss D, Appleboim A, Friedman N. 2019. Dynamics of Chromatin and Transcription during Transient Depletion of the RSC Chromatin Remodeling Complex. Cell Rep 26: 279–292 e275.

Kubik S, Bruzzone MJ, Jacquet P, Falcone JL, Rougemont J, Shore D. 2015. Nucleosome Stability Distinguishes Two Different Promoter Types at All Protein-Coding Genes in Yeast. Mol Cell 60: 422–434.

Kubik S, O’Duibhir E, de Jonge WJ, Mattarocci S, Albert B, Falcone JL, Bruzzone MJ, Holstege FCP, Shore D. 2018. Sequence-Directed Action of RSC Remodeler and General Regulatory Factors Modulates +1 Nucleosome Position to Facilitate Transcription. Mol Cell 71: 89–102 e105.

Kuras L, Struhl K. 1999. Binding of TBP to promoters in vivo is stimulated by activators and requires Pol II holoenzyme. Nature 399: 609–613.

Lai WKM, Pugh BF. 2017. Understanding nucleosome dynamics and their links to gene expression and DNA replication. Nat Rev Mol Cell Biol 18: 548–562.

Langmead B, Salzberg SL. 2012. Fast gapped-read alignment with Bowtie 2. Nat Methods 9: 357–359.

Li B, Howe L, Anderson S, Yates JR, 3rd, Workman JL. 2003. The Set2 histone methyltransferase functions through the phosphorylated carboxyl-terminal domain of RNA polymerase II. J Biol Chem 278: 8897–8903.

Malabat C, Feuerbach F, Ma L, Saveanu C, Jacquier A. 2015. Quality control of transcription start site selection by nonsense-mediated-mRNA decay. Elife 4.

Martens JA, Laprade L, Winston F. 2004. Intergenic transcription is required to repress the Saccharomyces cerevisiae SER3 gene. Nature 429: 571–574.

Mayer A, di Iulio J, Maleri S, Eser U, Vierstra J, Reynolds A, Sandstrom R, Stamatoyannopoulos JA, Churchman LS. 2015. Native elongating transcript sequencing reveals human transcriptional activity at nucleotide resolution. Cell 161: 541–554.

Mellor J, Woloszczuk R, Howe FS. 2016. The Interleaved Genome. Trends Genet 32: 57–71.

Miotto B, Ji Z, Struhl K. 2016. Selectivity of ORC binding sites and the relation to replication timing, fragile sites, and deletions in cancers. Proc Natl Acad Sci U S A 113: E4810–4819.

Murray SC, Haenni S, Howe FS, Fischl H, Chocian K, Nair A, Mellor J. 2015. Sense and antisense transcription are associated with distinct chromatin architectures across genes. Nucleic Acids Res 43: 7823–7837.

Neil H, Malabat C, d’Aubenton-Carafa Y, Xu Z, Steinmetz LM, Jacquier A. 2009. Widespread bidirectional promoters are the major source of cryptic transcripts in yeast. Nature 457: 1038–1042.

Nevers A, Doyen A, Malabat C, Neron B, Kergrohen T, Jacquier A, Badis G. 2018. Antisense transcriptional interference mediates condition-specific gene repression in budding yeast. Nucleic Acids Res 46: 6009–6025.

Nocetti N, Whitehouse I. 2016. Nucleosome repositioning underlies dynamic gene expression. Genes Dev 30: 660–672.

Nojima T, Gomes T, Grosso ARF, Kimura H, Dye MJ, Dhir S, Carmo-Fonseca M, Proudfoot NJ. 2015. Mammalian NET-Seq Reveals Genome-wide Nascent Transcription Coupled to RNA Processing. Cell 161: 526–540.

Porrua O, Libri D. 2015. Transcription termination and the control of the transcriptome: why, where and how to stop. Nat Rev Mol Cell Biol 16: 190–202.

Proudfoot NJ. 1986. Transcriptional interference and termination between duplicated alpha-globin gene constructs suggests a novel mechanism for gene regulation. Nature 322: 562–565.

Ramirez F, Ryan DP, Gruning B, Bhardwaj V, Kilpert F, Richter AS, Heyne S, Dundar F, Manke T. 2016. deepTools2: a next generation web server for deep-sequencing data analysis. Nucleic Acids Res 44: W160–165.

Rando OJ, Winston F. 2012. Chromatin and transcription in yeast. Genetics 190: 351–387.

Rhee HS, Pugh BF. 2012. Genome-wide structure and organization of eukaryotic pre-initiation complexes. Nature 483: 295–301.

Rundlett SE, Carmen AA, Kobayashi R, Bavykin S, Turner BM, Grunstein M. 1996. HDA1 and RPD3 are members of distinct yeast histone deacetylase complexes that regulate silencing and transcription. Proc Natl Acad Sci U S A 93: 14503–14508.

Sadeh R, Launer-Wachs R, Wandel H, Rahat A, Friedman N. 2016. Elucidating Combinatorial Chromatin States at Single-Nucleosome Resolution. Mol Cell 63: 1080–1088.

Sanchez-Gaya V, Casani-Galdon S, Ugidos M, Kuang Z, Mellor J, Conesa A, Tarazona S. 2018. Elucidating the Role of Chromatin State and Transcription Factors on the Regulation of the Yeast Metabolic Cycle: A Multi-Omic Integrative Approach. Front Genet 9: 578.

Schaughency P, Merran J, Corden JL. 2014. Genome-wide mapping of yeast RNA polymerase II termination. PLoS Genet 10: e1004632.

Schep AN, Buenrostro JD, Denny SK, Schwartz K, Sherlock G, Greenleaf WJ. 2015. Structured nucleosome fingerprints enable high-resolution mapping of chromatin architecture within regulatory regions. Genome Res 25: 1757–1770.

Schulz D, Schwalb B, Kiesel A, Baejen C, Torkler P, Gagneur J, Soeding J, Cramer P. 2013. Transcriptome surveillance by selective termination of noncoding RNA synthesis. Cell 155: 1075–1087.

Soudet J, Gill JK, Stutz F. 2018. Noncoding transcription influences the replication initiation program through chromatin regulation. Genome Res 28: 1882–1893.

Struhl K. 2007. Transcriptional noise and the fidelity of initiation by RNA polymerase II. Nat Struct Mol Biol 14: 103–105.

Tramantano M, Sun L, Au C, Labuz D, Liu Z, Chou M, Shen C, Luk E. 2016. Constitutive turnover of histone H2A.Z at yeast promoters requires the preinitiation complex. Elife 5.

van Nues R, Schweikert G, de Leau E, Selega A, Langford A, Franklin R, Iosub I, Wadsworth P, Sanguinetti G, Granneman S. 2017. Kinetic CRAC uncovers a role for Nab3 in determining gene expression profiles during stress. Nat Commun 8: 12.

van Werven FJ, Neuert G, Hendrick N, Lardenois A, Buratowski S, van Oudenaarden A, Primig M, Amon A. 2012. Transcription of two long noncoding RNAs mediates mating-type control of gametogenesis in budding yeast. Cell 150: 1170–1181.

Venkatesh S, Smolle M, Li H, Gogol MM, Saint M, Kumar S, Natarajan K, Workman JL. 2012. Set2 methylation of histone H3 lysine 36 suppresses histone exchange on transcribed genes. Nature 489: 452–455.

Venkatesh S, Workman JL. 2015. Histone exchange, chromatin structure and the regulation of transcription. Nat Rev Mol Cell Biol 16: 178–189.

Weiner A, Hsieh TH, Appleboim A, Chen HV, Rahat A, Amit I, Rando OJ, Friedman N. 2015. High-resolution chromatin dynamics during a yeast stress response. Mol Cell 58: 371–386.

Weiner A, Hughes A, Yassour M, Rando OJ, Friedman N. 2010. High-resolution nucleosome mapping reveals transcription-dependent promoter packaging. Genome Res 20: 90–100.

Wery M, Gautier C, Descrimes M, Yoda M, Vennin-Rendos H, Migeot V, Gautheret D, Hermand D, Morillon A. 2018. Native elongating transcript sequencing reveals global anti-correlation between sense and antisense nascent transcription in fission yeast. RNA 24: 196–208.

Xu Z, Wei W, Gagneur J, Perocchi F, Clauder-Munster S, Camblong J, Guffanti E, Stutz F, Huber W, Steinmetz LM. 2009. Bidirectional promoters generate pervasive transcription in yeast. Nature 457: 1033–1037.

Yen K, Vinayachandran V, Batta K, Koerber RT, Pugh BF. 2012. Genome-wide nucleosome specificity and directionality of chromatin remodelers. Cell 149: 1461–1473.

Zentner GE, Henikoff S. 2013. Regulation of nucleosome dynamics by histone modifications. Nat Struct Mol Biol 20: 259–266.

Zentner GE, Kasinathan S, Xin B, Rohs R, Henikoff S. 2015. ChEC-seq kinetics discriminates transcription factor binding sites by DNA sequence and shape in vivo. Nat Commun 6: 8733.

